# Multi-omics comparison of malignant and normal uveal melanocytes reveals novel molecular features of uveal melanoma

**DOI:** 10.1101/2022.03.11.483767

**Authors:** David Gentien, Elnaz Saberi-Ansari, Nicolas Servant, Ariane Jolly, Pierre de la Grange, Fariba Némati, Géraldine Liot, Simon Saule, Aurélie Teissandier, Deborah Bourchis, Elodie Girard, Jennifer Wong, Julien Masliah Planchon, Manuel Rodrigues, Laure Villoing Gaudé, Cécile Reyes, Matéo Bazire, Thomas Chenegros, Mylene Bohec, Sylvain Baulande, Radhia M’kacher, Eric Jeandidier, André Nicolas, Didier Decaudin, Nathalie Cassoux, Sophie Piperno-Neumann, Marc-Henri Stern, Johan Harmen Gibcus, Job Dekker, Edith Heard, Sergio Roman-Roman, Joshua J. Waterfall

## Abstract

Uveal Melanoma (UM) is a rare cancer resulting from transformation of melanocytes residing in the uveal tract. Integrative analysis has identified four molecular and clinical subsets in UM. To improve our molecular understanding of UM we performed extensive multi-omics characterization comparing two aggressive UM patient-derived xenograft models with normal choroidal melanocytes, including DNA optical mapping, specific histone modifications, and DNA topology analysis by Hi-C. Our gene expression and cytogenetic analyses suggest that genomic instability is a hallmark of UM. We also identified a recurrent deletion in the *BAP1* promoter resulting in loss of expression and associated with high risk of metastases in UM patients. Hi-C revealed chromatin topology changes associated with up-regulation of *PRAME*, an independent prognostic biomarker in UM and potential therapeutic target. Our findings illustrate how multi-omics approaches can improve the understanding of tumorigenesis and reveals two novel mechanisms of gene expression dysregulation in UM.

## Introduction

Uveal Melanoma (UM) is a rare cancer (5-7 cases per million per year) which affects mainly adults and represents 5% of all melanoma [1]. UM results from the malignant transformation of melanocytes of the uveal tract of the eye, which comprises the iris, the ciliary body, and choroidal membrane [2]. UM primary tumors are well controlled by surgery and/or radiotherapy but more than 30% of the patients develop metastases, mainly in the liver, with a very poor prognosis. Improvement in the understanding of aggressive UM is essential for identifying efficient new therapeutic approaches.

The vast majority of UM display activating mutations of *GNAQ* [3] or its paralog *GNA11* [4], their upstream activator *CYSLTR2* [5] or downstream effector *PLCB4* [6]. These mutually exclusive Gα/q-related mutations present in 98% of UM are recognized as a primary event of UM oncogenesis [7] and lead to the activation of Gα/q signaling pathway [8], [9]. Mutations in *BAP1, EIF1AX, SF3B1,* and *SRSF2,* [10]–[13] were identified as a secondary mutational event necessary for malignant transformation. Mutations in *BAP1, SF3B1,* and *EIF1AX* (so called BSE events) are associated with distinct delays in the appearance of metastasis with the shortest delay associated with *BAP1* [14].

Over the last two decades, a number of recurrent chromosomal abnormalities have been identified in UM, including monosomy 3 (M3), gains of 6p and 8q, as well as loss of 6q and 8p. These abnormalities are associated with adverse clinical outcome and are currently used for clinical prognosis [15]–[18]. Monosomy 3 and gain of chromosome 8 correlate with an intermediate risk of metastasis and the highest risk of metastasis is associated with combined M3 and gain of 8q [16], [18]. Integrative analysis including copy number variations, DNA methylation, recurrent mutations, and gene expression profiles has identified four molecular and clinical subsets in UM [18].

To improve our understanding of tumor oncogenesis, we performed extensive multi-omic and FISH characterization of two aggressive UM PDX with distinct mutational and chromosomal rearrangement patterns as well as short term culture of normal choroidal melanocytes (NM) for comparison. In addition to investigating somatic DNA alterations and performing RNA sequencing and DNA topology analysis, we also performed whole genome DNA methylation sequencing and chromatin immunoprecipitation (ChIP-seq) of histone marks associated with activating (H3K4me3), repressing (H2A119Ub, H3K27me3) or enhancing (H3K27Ac) gene expression. These complementary analyses improved the characterization of regulated genes and pathways in aggressive UM.

## Results

### Samples studied

We sorted UM cells from two aggressive PDX models, MP41 and MP46 [19]–[21], to obtain both a pure tumor population and a sufficient number of cells [22]. The MP41 model was generated from enucleation of a UM occurring in a 50-year-old female patient who had a metastasis 31 months after initial diagnosis and who died 43 months after diagnosis from multiple metastases (including bones, lung, and subcutaneous lesions). The MP46 model was established from enucleation of a tumor occurring in a 69-year-old male patient. This patient developed a liver metastasis 6 months after diagnosis of the primary tumor, and died 7 months from initial diagnosis (Figure 1A). These two aggressive models harbor canonical activating mutations in *GNAQ/11* genes and share 8q and 6p gains. MP46 displays monosomy of chromosome 3 and is deficient in BAP1 by immunohistochemistry (IHC) (Figure 1B), even though no *BAP1* mutations were identified by Sanger sequencing. MP41 is BAP1 proficient by IHC (Figure 1B) and no mutations were identified in either *BAP1, SF3B1 or EIF1AX*.

**Figure 1:**
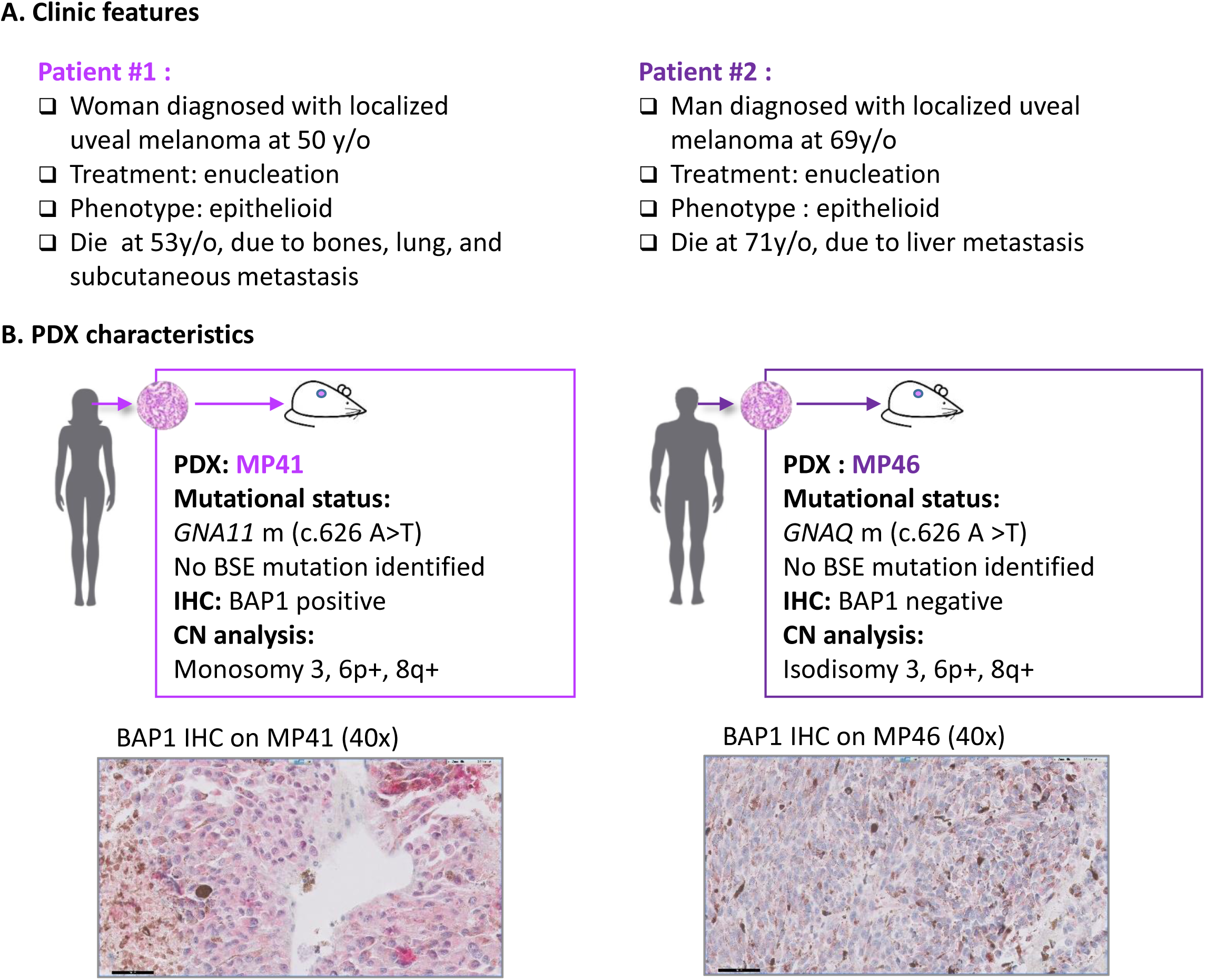
Principal characteristics of MP41 and MP46 PDXs established from aggressive uveal melanomas. A) Clinical characteristics of UM cases. B) Main molecular characteristics of corresponding patient derived xenograft models established and characterized previously [21]. Mutational status was assessed with Sanger sequencing (*GNAQ/GNA11, BAP1, SF3B1, EIF1AX*), with Cytoscan HD microarrays for copy number analysis and BAP1 immunohistochemistry from FFPE tissue section.

### Whole genome sequencing and copy number analysis confirmed MP41 and MP46 as high risk UM

First, MP41 and MP46 were subjected to whole genome sequencing (WGS) to perform single nucleotide variant (SNV) and copy number analyses. To facilitate the identification of somatic alterations, WGS was also conducted on matched healthy tissue adjacent to original primary tumors.

Somatic point mutation analysis revealed less than one somatic mutation per Mb (0.42 and 0.37 SNV/Mb in MP41 and MP46 respectively) as observed previously in UM [18]. Based on Cancer Genome Interpreter [23] and VarSome [24] classification, a unique known driver mutation associated to a pathogenic role was retrieved in both MP41 (*GNA11* c.626A>T, allele frequency (AF):43%) and MP46 (*GNAQ* c.626A>T, AF: 43%). Supplementary Tables 1 and 2 list all the SNVs labelled as passenger mutations, having a moderate to high impact on amino acid sequence, or affecting ncRNAs for MP41 and MP46, respectively. Additional mutations identified in MP41 included a premature stop codon in *KMT2C* (KMT2C:[p.Tyr987*]), a known driver mutation and 13 passenger mutations (based on oncodrive MUT algorithm). In MP46, we identified 21 passenger mutations in 19 genes of which 8 were predicted to be pathogenic.

The comprehensive TCGA UM study distinguished four copy number subtypes that had diverse aneuploid events and divided D3-UM and M3-UM into two subgroups, based on somatic copy number alterations [18]. Somatic copy number alterations as losses (L) and gains (G) identified from WGS of MP41 and MP46 models include notably for MP41: M3, G6p, L6q, L8p, G8q, and for MP46: Isodisomy 3, G6p, L8p, G8q as published [21] (Supplementary Table 3). Based on the TCGA copy number subtypes of uveal melanomas, MP41 and MP46 were classified in group 2 and group 4 respectively [18]. The classification of MP46 is consistent with the inclusion of BAP1-deficient tumors in group 4. Although TCGA group 2 is enriched in *SF3B1*-mutated UM, no *SF3B1* or *SRSF2* mutation and no *SF3B1* splicing patterns were observed in MP41 [13].

Overall, WGS analysis of MP41 and MP46 models confirmed the presence of a unique oncogenic driver mutation in the Gαq pathway, the presence of M3 and G8q, and an association of TCGA copy number group 2 for MP41 and group 4 for MP46.

### Gene expression analysis reveals upregulation of well-known genes and highlights DNA repair pathways

We performed a gene expression analysis to compare the transcriptome of UM models to normal choroidal melanocytes. The RNAseq dataset was composed of five normal melanocytes cell lines (NM) including a technical replicate, four MP41 biological replicates and three MP46 biological replicates. Using unsupervised principal components analysis (PCA) and hierarchical clustering of all differentially expressed genes compared to NM controls (log2 fold change (FC)>1.5, p-value ≤ 5%) we found a high reproducibility of the replicates and clear separation between UM models and NM (Figure 2A-B).

**Figure 2:**
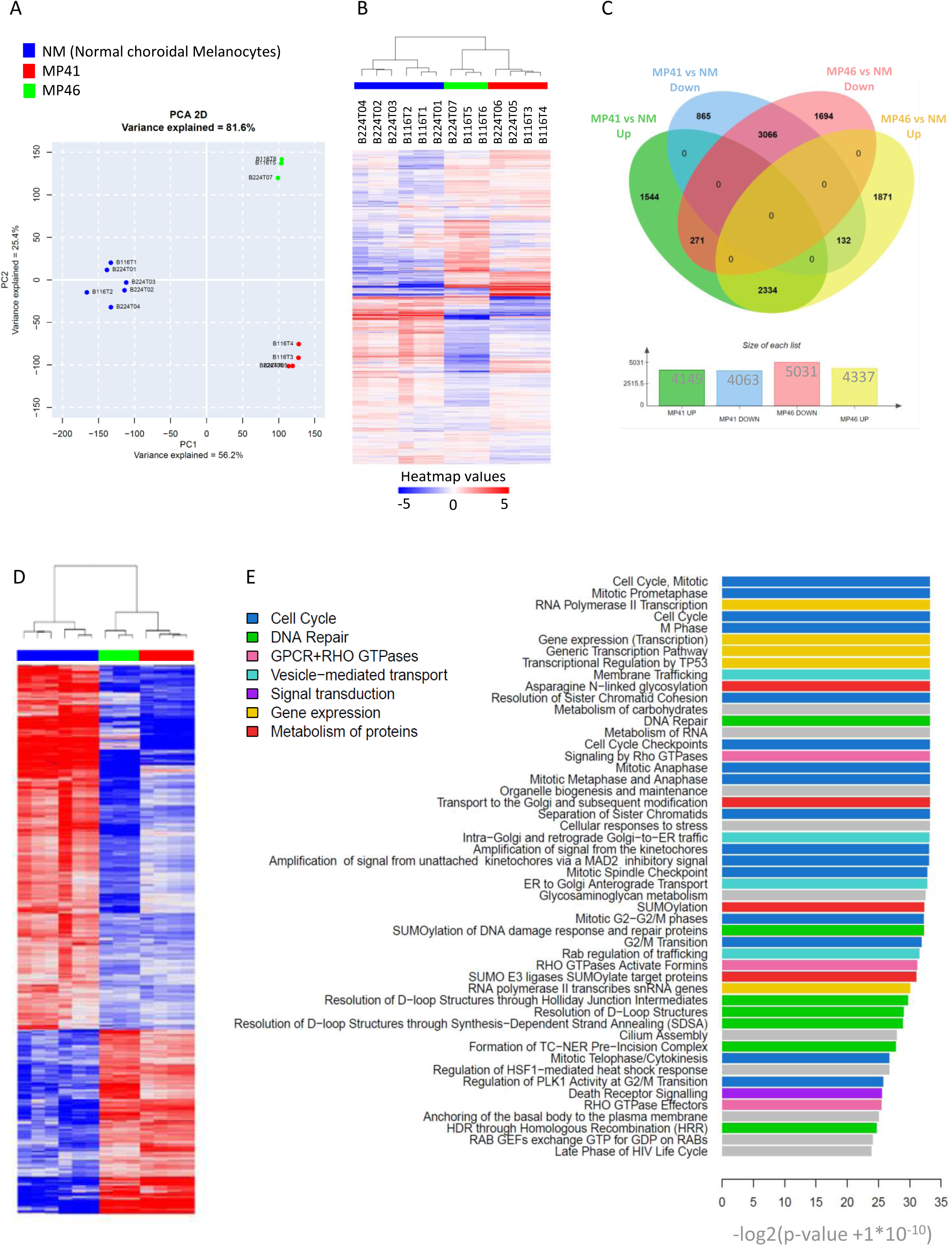
Gene expression global overview. A) Principal component analysis of RNASeq of 6 normal choroidal melanocyte samples (blue), 4 preparations of MP41 UM cells (red) and 3 preparations of MP46 UM cells (green). B) Hierarchical clustering of same profiles. C) Differential gene expression analysis of MP41 vs NM and MP46 vs NM to identify genes with a Log2 fold change greater than 1.5 and a p-value lower than 0.05. D) Heatmap of commonly regulated genes in MP41 vs NM and MP46 vs NM. E) 50 most highly regulated pathways by Reactome analysis of commonly regulated genes listed according the significance (-log 2 (p-value +1x10^-10^)).

To identify consistently differentially expressed genes (DEG) in aggressive UM, we compared each PDX to the NM and then compared the resulting gene lists (Figure 2C, Supplementary Table 4). For MP41, a total of 8,212 DEG were identified (4,149 up and 4,063 down). Among the 9,368 genes identified in MP46, 4,337 were over-expressed and 5,031 were under-expressed. The overlapping 3,066 downregulated genes and 2,334 upregulated genes between MP41 and MP46 (Figure 2D, Supplementary Figure 1) were subjected to further analysis.

Cancer testis antigens were significantly enriched among the consistently over-expressed genes in MP41 and MP46, with *PRAME* [25] as the highest cancer testis antigen expressed in both MP41 and MP46 (log2 FC: ∼12). *PLCB4* and *RASGRP3*, two key genes in UM oncogenesis were among the top 50-upregulated genes. A small percentage of UM patients display activating mutations in the PKC regulator *PLCB4* which are mutually exclusive with *GNAQ/GNA11/CYSLTR2* mutations [6]. The overexpression of *PLCB4* suggests a potential contribution to the activation of Gαq pathway in the absence of *PLCB4* activating mutations.

*RASGRP3* has been shown to mediate MAPK pathway activation in UM [8], [9]. We identified an additional GPCR downstream pathway gene, *RAPGEF4*, as significantly upregulated in both models.

Ranking the consistent DEGs by fold change indicated that the top 50 most upregulated genes in MP41 and MP46 were quite similar (Table 1A-B). Several non-coding RNAs including HAGLR and TRPM2-AS, which have been previously reported to participate in oncogenesis [26]–[30], were found over-expressed. This suggests that other consistently over-expressed genes may also play functional roles in UM. The overlap between differentially downregulated genes in MP41 and MP46 was less pronounced (Table 1C and D). Only eight of the top 50 most downregulated genes were shared between models. Consistent with the IHC result, BAP1 was the most downregulated gene in MP46.

**Table 1:**
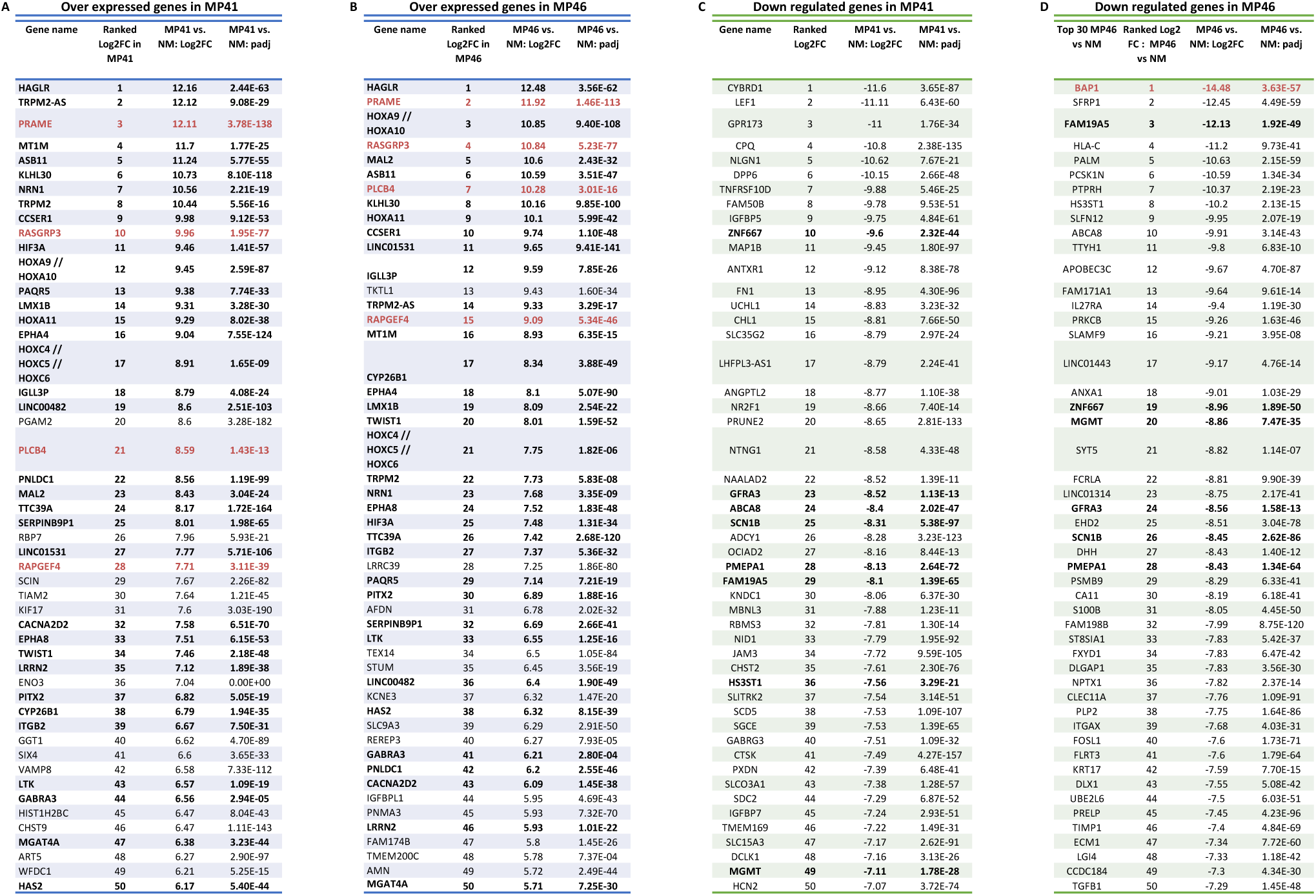
Top 50 of regulated genes in UM models vs NM. Gene are ranked according log 2 fold changes. A and B. Top 50 of over-expressed genes in MP41 and MP46 respectively. C and D Top 50 of down under-expressed genes in MP41 and MP46 respectively. Genes written in bold are regulated with the same variation in both comparisons. Genes written in red were previously identified in UM studies.

We identified 101 cytobands that contained DEG (Supplementary Figure 1) most frequently located on 8q (18%). Up-regulated genes were significantly associated with copy number increases on chromosomes 8q, 1q, 21q in MP41 and MP46. We also found upregulated genes on cytobands from 5q and 4q, which have normal copy numbers (CN) in MP41 and MP46, and on cytobands from 2p/2q and 7q that were gained only in MP46. Down-regulated genes were associated with loss of 1p, 3p, 8p and 16q in MP41 and MP46. Yet, we also found significantly downregulated genes on 9q, 10q, 19p/q, lost only in MP41, and on 12q and 17q that were lost only in MP41 and MP46 respectively.

We next used Reactome [31] analysis to identify enriched pathways in the DEG. The top 50 dysregulated pathways are shown in Figure 2E. These pathways include proliferation-related pathways (cell cycle, mitosis, checkpoints) as well as chromatin maintenance and DNA repair pathways (DNA double strand break repair, Fanconi anemia) (p <1.5x10^-4^, Supplementary Figures 2). DNA damage and repair (DDR) pathways were found to be enriched, and within DDR pathways, the Homologous Recombination (HR, Supplementary Figure 2B) pathway was the most significantly enriched (p: 5.99x10^-9^) based on the enrichment of DNA damage sensors and repair enzymes genes. Additionally thirteen genes from within the Fanconi Anemia pathway (FA, Supplementary Figure 2C) were consistently enriched after comparing UM to NM (min p-value: 2.94x10^-4^).

In summary, these analyses revealed a list of DEG from two UM models compared to NMs with notably the over-expression of two GNA11/GNAQ pathway downstream genes: RAPGEF4 and PLCB4. Interestingly, DNA damage repair pathways were significantly enriched and PRAME, a marker of aggressiveness in UM, was among the most upregulated genes in both MP41 and MP46.

### Optical mapping and FISH analyses reveal major chromosomal aberrations

To further investigate genomic aberrations related to UM, we performed optical mapping with the Bionano platform [32], [33] as well as telomere and centromere staining followed by M-FISH (TC+M-FISH [34]) on MP41 and MP46. The optical mapping achieved 500bp resolution and a minimum coverage of 97X per sample, which revealed long-range DNA alterations including translocations, insertions, duplications, and small deletions in both models (Figure 3A, B and C). In MP41, optical mapping revealed both intra- and inter-chromosomal translocations (t(19;19) t(1;12) and t(6;8)). In MP46, intra-chromosomal translocations for chr19 and the inter-chromosomal translocation t(1;22) were identified. Structural variants (SVs) including deletions, insertions, and duplications were also identified and summarized in Figure 3C.

**Figure 3:**
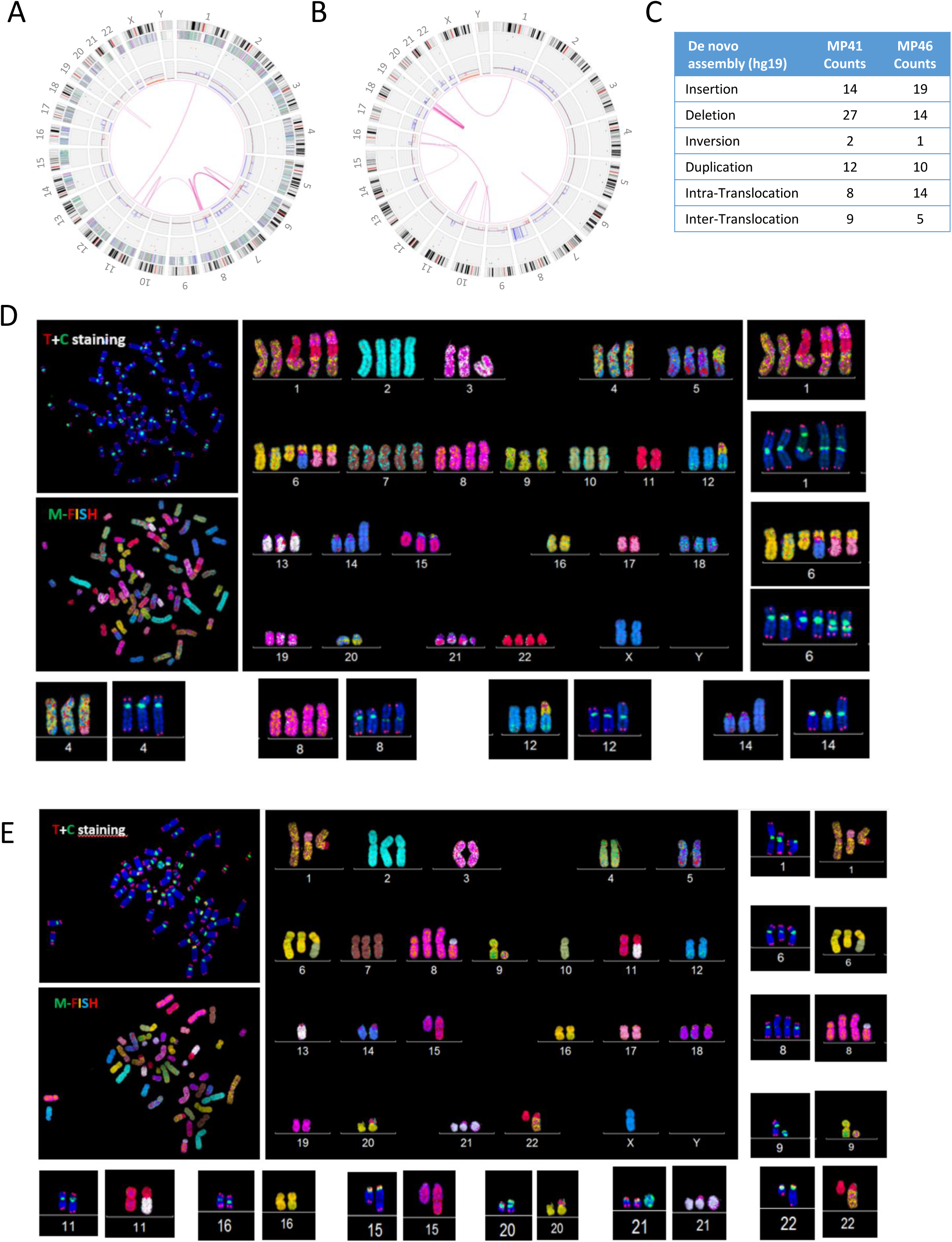
DNA optical mapping and FISH analysis of MP41 and MP46. Circos plot of aberrations in A) MP41 and B) MP46. From the central to the periphery of the circos plot: whole genome view summarizes intra and inter chromosomal translocations (pink lines), copy number gains and losses listed on the first internal layer of the circos, and SVs (insertion, deletion inversion and duplication) are labelled as colored dots in the intermediate layer of the circos. Gene density, cytobands and chromosomes are composing the outer layers of the circos. C) Number of insertions, deletions, inversions and duplications and inter and inter translocations are detailed for MP41 and MP46 defined by Bionano optical mapping. Telomere and M-FISH from analysis of D) MP41 and E) MP46 are derived from two different FISH analyses. Upper left panels show telomere (red signal) and centromere (green signal) staining and counter labelled with DAPI (blue). Lower left panels show M-FISH analysis. Main panels correspond to the karyotype view.

TC+M-FISH revealed a hyper-triploid genome for MP41 (Figure 3D) with dicentric chromosomes (dic(14;16), i(8q), dic(1;11;8)). We also identified four telomeric-related losses dic(1;11;6;8), dic(1;11), dic(6;8;14) and dic(6;8;17), and one interstitial telomeric sequence dic(6;8;17) (Supplementary Figure 3). Chromosomal end-to-end fusions are often associated with dicentric chromosomes and aberrant chromosomal structures. MP46 also displays a complex karyotype (Figure 3E): a hyper-diploid genome with multiple dicentric chromosomes: dic(1;17), dic(6;10), dic(8;21) and dic(13;22), dic(16;20); dic(20;22). Two translocations were also identified in MP46: t(1;22) and der(15)t(11;15). In addition to chromosome structural alterations, we repeatedly found ring chromosomes derived from chromosomes 1, 8, and 11 in MP41 and from chromosomes 9 and 21 in MP46 (Supplementary Figure 3E).

We next analyzed the RNAseq data for potential fusions using multiple algorithms and confirmed with RT-PCR and Sanger sequencing. In most cases, SVs were directly associated with the presence of fusion RNAs. For example, an insertion-duplication on 12q24 in both MP41 and in MP46, led to the fusion of MAPKAPK5-ACAD10 in MP41 and KDM2B-RHOF in MP46. Both events occurred within the replication fragile site FRA12E [35] which has also been associated with germline structural polymorphisms [36]. In MP46, a t(1;22) translocation between CABIN1 and MPRS21 led to the fusion RNA CABIN1-MPRS21.

Interestingly, a complex translocation t(6;8) in MP41 resulted in two fusion RNAs: *GPAT4-NCOA7* and *POMK-RSPO3*. From FISH, we determined that the fusion was located on derivative chromosomes 6: der(6)t(6;8); dic(der(6)t(6;8);14); dic(der(6)t(6;8);16); dic(der(6)t(6;8);17), and derivative chromosomes 8: der(8)t(6;8), ider(8)(q10)t(6;8)x2, dic(1;11;8;6)x2, and on a ring chromosome r(dic1;11;8;6). In MP41, we specifically probed *NCOA7* and *RSPO3* (both on chromosome 6), as well as *GPAT4*, and *POMK* (both on chromosome 8) (Supplementary Figure 4A). We found that, contrary to dicentric chromosome 6, which was labelled with all four probes, normal chromosomes 6 only stained with *NCOA7* and *RSPO3* probes (Supplementary Figure 4B, C and D). On chromosome 8 we could stain the native genes *GPAT4* and *POMK* as well as chromosome 6 derived *NCOA7* (but not *RSPO3*) (Supplementary Figure 4E). Isochromosome 8 was only labelled with *GPAT4* and *NCOA7*, as dic(1;11;8;6) (Supplementary Figure 4E and F). A potential model for the generation of these complex patterns is in Supplementary Figure 4G.

We next investigated structural genomic aberrations in additional models of aggressive UM (Mel202, MM66, OMM1, and OMM2.3) through optical mapping and FISH analyses (Supplementary Figure 5A). Major SVs were detected in all tested UM models (Supplementary Figure 5B). Telomere aberrations were present in all tested UM cell lines (MP41, MP46, Mel202, MM66, OMM1 and OMM2.3 [37], [38]) and absent in normal controls (Supplementary Figure 5C). Translocations and dicentric chromosomes were detected in the cellular models (Mel202, MM66; OMM1; OMM2.3, Supplementary Figure 5D-G). Derivative chromosomes were also associated with complex SVs identified with optical mapping.

To summarize, high resolution DNA optical mapping combined with TC+M-FISH shows that high levels of genomic instability are a recurrent pattern in diverse aggressive UM models.

### DNA methylation analysis reveals differences in CpG island (CGI) patterns and identifies BAP1 promoter deletion

To characterize DNA methylation patterns, OxBS sequencing (Cambridge Epigenetix) was performed on both UM and NM. Oxidative bisulfite sequencing [39] was found to be more robust in our samples, particularly for the normal melanocytes, most probably due to the abundance of melanin.

First, a random forest analysis placed MP41 and MP46 in the TCGA methylation group 2 (corresponding to BAP1 proficient) and group 4 (BAP1 deficient) respectively. We next categorized methylation levels with respect to the following genomic localizations: CGI promoters, non-CGI promoters, exons, introns, intergenic regions and repeat elements (identified by RepeatMasker annotations [40]). As illustrated in Figure 4A, whereas CGI-promoters are generally equally demethylated in UM samples and NM, tumor samples are globally less methylated in non-CGI promoters as compared to normal samples across all genomic localization categories [41], [42].

**Figure 4:**
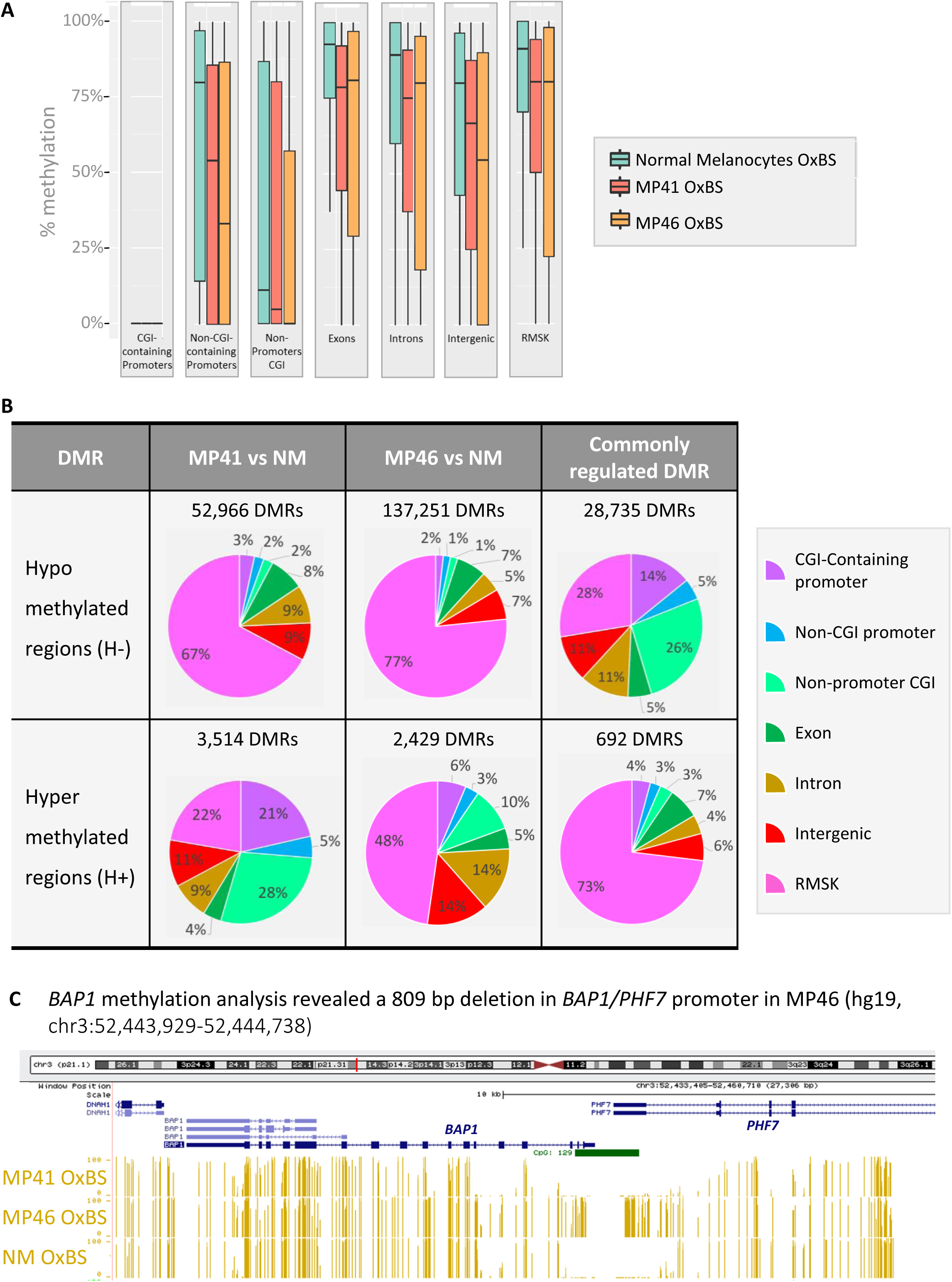
Whole genome DNA methylation analysis of UM models and normal melanocytes. A) DNA methylation levels based on oxidative bisulfite DNA treatment followed by whole genome sequencing, is shown at CGI promoters, non- CGI promoters, non-promoters CGI, exons, introns, intergenic regions, and on repeats, in normal melanocytes, MP41 and MP46. B) Differentially Methylated Regions (DMRs) as hypo and hyper DMRs (H- and H+) in 300kb window in MP41 vs NM, MP46 vs NM. Commonly regulated DMR correspond to DMR identified in MP41 vs NM and MP46 vs NM comparisons. DMR were considered as commonly regulated where sharing the same variation (H- or H+) and when their coordinates were identical or overlapping. C) Percentage of CpG methylation in MP41, MP46 and NM in *BAP1* locus through UCSC Genome Browser is represented in yellow. CpG Island 129 overlaps the *BAP1/PHF7* promoter.

Next, we identified differentially methylated regions (DMR) in each PDX compared to NM (Figure 4B). Most DMRs are hypomethylated (H-) in agreement with global methylation patterns (Figure 4A). We focused our analysis on DMRs which are consistent in MP41 and MP46. As expected most shared DMR are H- (28,735 / 29,427 - 98%). Notably, half of MP41 DMRs are shared with MP46, but MP46 shares less than a third of its DMR with MP41 (H-: 21%, H+ :28%). Interestingly, while H- genomic localizations are strongly enriched at repeat elements in each model independently, the shared H- are strongly depleted at repeat elements in each model independently, the shared H- are strongly depleted for this elements (Figure 4B). Genomic localizations of H+ DMRs are differently distributed in MP41 and MP46 with the largest differences occurring in repetitive elements and CpG islands. The localization of the 692-shared H+ is primarily in repeat elements and a subset is located in both CGI- and non-CGI promoters (Figure 4B).

As mentioned before, MP46 clusters with the DNA methylation group 4 of TCGA [18], which includes BAP1- deficient and monosomy 3 UM tumors. We analyzed the status of *BAP1* promoter methylation, because we did not find *BAP1* mutations in MP46 [21] despite the absence of BAP1 protein. Although MP41 displays a hypo-methylated promoter similar to NM, consistent with expression levels (Supplementary Figure 6 A), MP46 shows a specific hyper-methylation pattern in the promoter (CpG129, UCSC genome browser (hg19; chr3:52,443,678-52,445,104) that co-localized with the boundary of a 809bp deletion identified in the whole genome OxBS data (Figure. 4C). The *BAP1* promoter is a bidirectional promoter shared between *BAP1* and *PHF7*. In MP46, the *BAP1* promoter was deleted, and deletion boundaries were hyper-methylated.

Furthermore, both *BAP1* and *PHF7* were not expressed in MP46 (Supplementary Figure 6A and B). This large deletion was also confirmed in our DNA labeling, optical mapping and WGS data.

BAP1 promoter deletion has not been described in the 1346 ClinVar records or in the Cosmic database. To investigate if promoter deletion explains other cases of BAP1 deficiency, a targeted NGS approach based on tiling amplicon-sequencing covering BAP1 was performed on 53 tumor samples (Supplementary Figure 6C). We identified two additional cases with a similar deletion in the *BAP1* promoter (Supplementary Figure 6D), for which immunohistochemistry confirmed the absence of BAP1 expression (Supplementary Figure 6E). A recent UM case has also been identified internally harboring a deletion of *BAP1/PHF7* promoter, wider than that observed in MP46 (2.2kb). In the tumor, BAP1 could not be detected by IHC and the promoter deletion was confirmed by long range PCR (Supplementary Figure 7). This recent UM case was analyzed as part of the French initiative *France Medecine Genomique 2025* and made possible by the SeqOIA platform (https://pfmg2025.aviesan.fr/en/).

Next, we analyzed the methylation status of the consistent DEG in MP41 and MP46. Most of the DEG did not a have a significant methylation switch, since only 5% of DEG displaying hypo or hyper-methylation of their promoters. However, several of the most differentially expressed genes, such as RASGRP3 and PRAME discussed above, were part of this minority.

### DNA topology analysis reveals stable compartments and TADs containing most differentially expressed genes

The spatial organization of melanocyte genomes and in particular physical interactions may contribute to the regulation of gene expression during transformation. We performed chromosome conformation capture (Hi-C) to elucidate if gene expression changes are associated with chromatin organization and DNA folding in MP41 and MP46 (Figure 5A). To account for background from copy number alterations (Supplementary figure 8), we first compared multiple computational approaches for normalization. We found that CAIC was less sensitive to copy number variations than ICE and LOIC (Supplementary Figure 9 and 10) similar to previously analyzed breast cancer cell lines [43]. Genome folding presents itself at multiple length scales: chromosome territories contain physically separated euchromatic and heterochromatic regions known as A and B compartments [44] and topologically associated domains (TADs) that result from loop extrusion [45]–[48].

**Figure 5:**
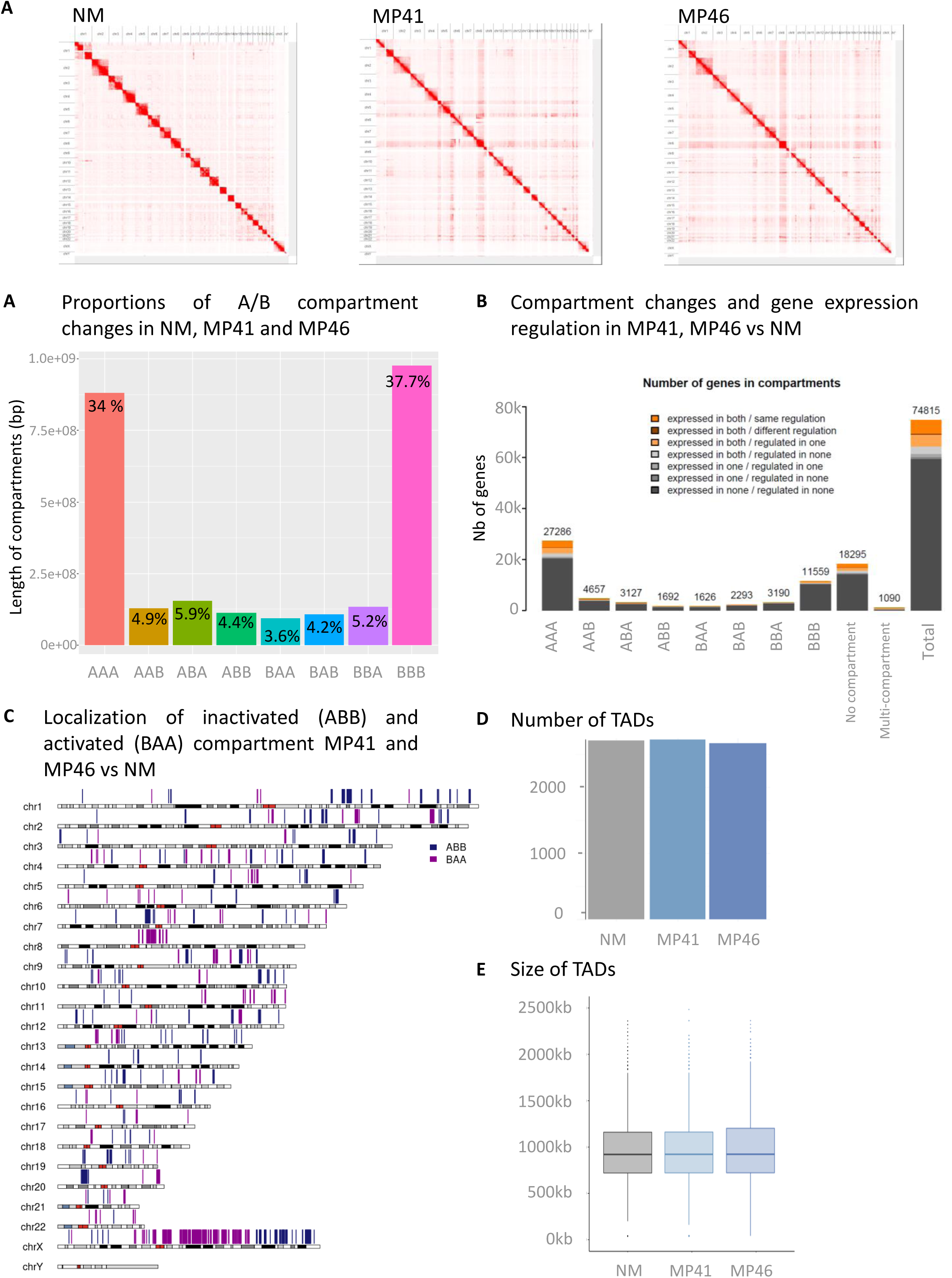
Compartments and TADs in NM and UM. A) Contacts maps derived from in situ Hi-C at the whole genome level for NM, MP41 and MP46. B) Histogram of compartment changes in NM, MP41 and MP46. A- and B-compartments identified at a 250kb resolution. C) Localization of inactivated (ABB) and activated (BAA) compartment in MP41, MP46 vs NM on a whole genome view. D) Integration of compartment changes and gene expression between NM, MP41 and MP46. E) Number and F) size of TADs in NM, MP41 and MP46.

First, we performed a compartment analysis at 250 Kb resolution [44], in NM, MP41 and MP46 (and refer to each window with a three letter code for its compartment status in NM, MP41 and MP46 respectively). Most compartments (∼72%) shared the same status between our 3 models: A compartments (labelled as ”AAA”, 34.04%) and B compartments (BBB: 37.72%) (Figure 5B). Of note, the CAIC normalization allowed equal detection of A/B compartments irrespective of copy number status (Supplementary Figure 10C). The other compartment patterns are roughly frequent, accounting for 3.5-6% of windows (Figure 5B). A karyotype view was used to illustrate the position of compartment assignment changes in NM and UM models (Figure 5C). We further analyzed whether changes in compartment status overlapped with gene content and gene expression (Figure 5D). Most of the DEG were located in the constitutive A compartments (AAA), this enrichment was not significant when corrected for the total number of genes and the number of genes per compartment. The ABB and BAA compartments contained 159 DEGs (same regulation, 96 in ABB and 63 in BAA), but this was not statistically significant (Supplementary Table 5). Second, TADs were analyzed in both UM models using insulation score analysis in 40kb bins [49]. No significant differences in the number or size of TADs were observed between UM and NM (Figure 5E, 5F).

In summary, no differences in TAD structures were found and most differentially expressed genes (83%) were found in compartments that did not change status from NM to UM (Figure 4C). However, 159 DEG (∼3%) were associated with changes in compartment status: 63 DEG belong to active compartments in MP41 and MP46, and 96 DEG belong to inactive compartments, indicating that differences in topology could underly differences in gene expression.

### Chromatin topology and histone marks changes are associated with up-regulation of *PRAME*

To further enrich our understanding of chromatin organization and gene expression regulation in NM, MP41 and MP46, chromatin immunoprecipitation (ChIP) and sequencing analysis were carried against the active epigenetic mark H3K4me3, and repressive marks H2AUb and H3K27me3. Additionally, in MP41 and MP46, we used H3K27Ac to find active enhancers and CTCF to sudy cohesin-mediated loop extrusion [50].

As depicted in Figure 5D, 371 active compartments were identified specifically in tumor setting (BAA) containing 63 consistently DEG (37 higher and 26 lower) in MP41 and MP46 versus NM. Among the 37- upregulated genes in activated compartments, *PRAME* and *ZNF280A* were found enriched in H3K4me3. Also, 33 genes (including *PRAME, ZNF280A/B, EZH2*) display H3K27Ac peaks, 2 genes lost H3K27me3 marks (*PITX2* and *COL4A5*) and 4 were demethylated in their promoter. Among the 26 downregulated genes, none were enriched in H3K27me3 marks in both UM models, only one gene (*ZC4H2*) lost H3K4me3, 20 genes contain H3K27Ac marks, and no gene displays a hyper methylated promoter.

Upregulated genes associated with activated compartments include *EZH2*, *EPHA4*, and *PRAME*. Among the major regulated genes, *PRAME* is associated with a particularly high fold change (log2 Fold Change ∼12.1 in MP41 and 11.9 in MP46 vs NM). *PRAME* upregulation was identified through a standard differential gene expression analysis RNAseq (Easana, Figure 6A). Although an absence of PRAME expression is observed in our NM, a huge number of *PRAME* counts are present in MP41 and MP46.

**Figure 6:**
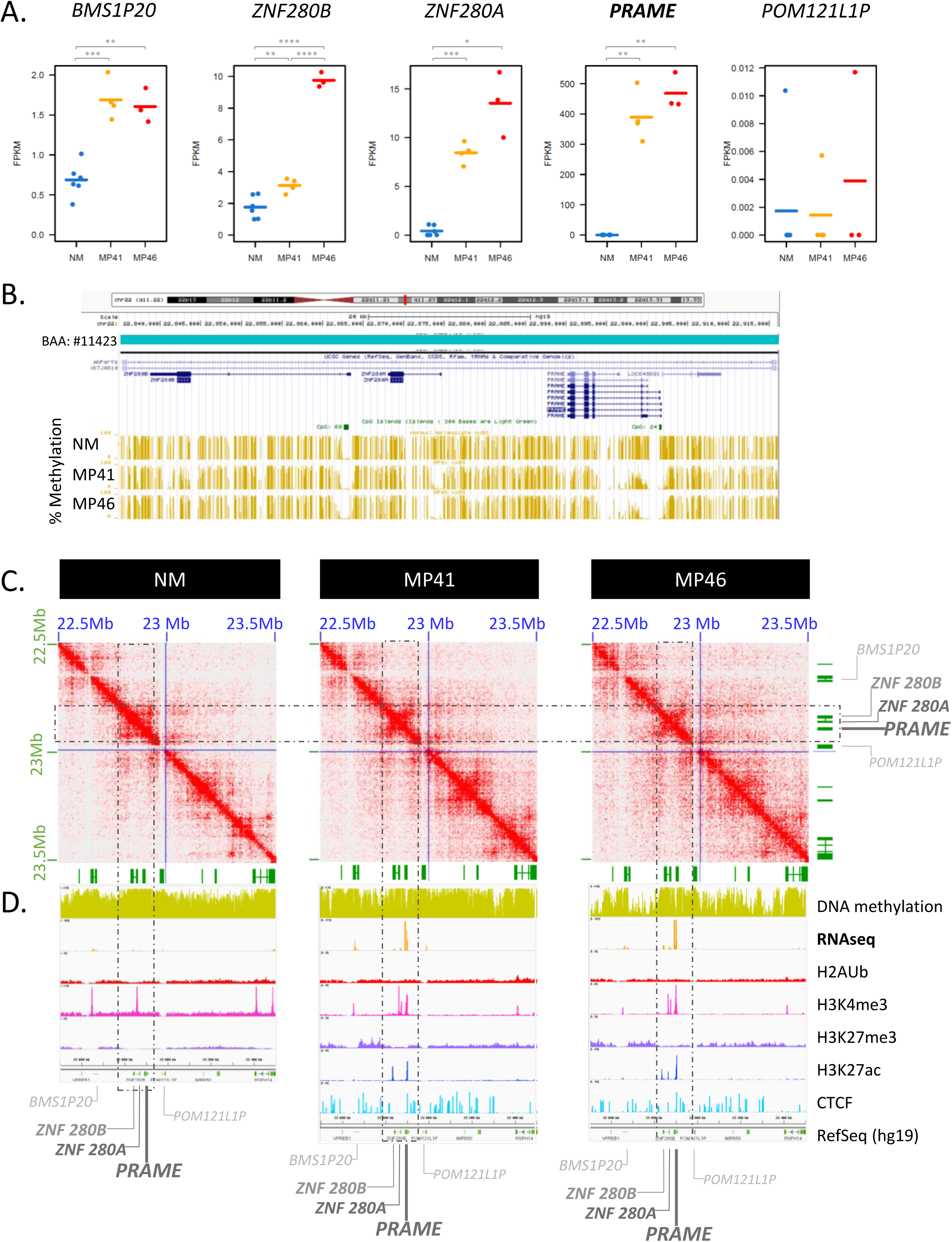
Multiomics analysis of *PRAME* locus. A) Gene expression of *PRAME* and its neighbors as *BMS1P20, ZNF280B, ZNF280A* upstream genes and POM121LP downstream. RNAseq data of NM, MP41 and MP46 replicates (FPKM). B) UCSC Genome Browser view (hg19) of percentage of DNA methylation (golden bars). C) DNA contacts maps of NM, MP41 and MP46 at 5kb resolution in *PRAME* TAD (blue square). D) UCSC Genome Browser view of the *PRAME* locus showing DNA methylation, RNAseq (log2), H2AUb, H3K4me3, H3K27me3, H3K27Ac, CTCF and RefSeq genes.

The *PRAME* gene is situated on 22q11.22 (hg19 chr22:22,890,123-22,900,022) between tandem Zinc finger proteins *ZNF280A* and *ZNF280B* previously identified as Suppressor of Hairy Wing (*SUHW1/ZNF280A* and *SUHW2/ZNF280B*) downstream of *PRAME* and a gene encoding a putative membrane glycoprotein (*POM121L1*) upstream of *PRAME* as illustrated on Figure 6D. In this locus, only *ZNF280A*, *ZNF280B* and *PRAME* are upregulated in a compartment being activated in our UM models compared to NM (Figure 6A). We also observed hyper methylation of the *PRAME* promoter in normal melanocytes and an hypo- methylation in MP41 and MP46 tumor models (Figure 6B), which correlates with the activation of *PRAME* expression in UM (Figure 6A and 6D) [51]. The *PRAME* promoter methylation status has previously been described in UM as a marker of aggressiveness [52].

Our analysis of histone marks at the *PRAME* locus in MP41 and MP46, revealed the presence of active transcription histone marks (H3K4me3, H3K27Ac), and the absence of repressed transcription histone marks such as H3K27me3. In MP46 an additional H3K27Ac peak is observed in *ZNF280A*, which potentially also contributes to *ZNF280A* expression. In NM, H3K4me3 and H3K27me3 peaks were not observed on the *PRAME* promoter (Figure 6D).

Chromatin conformation analysis at the *PRAME* locus revealed a different pattern in NM compared to UM MP41 and MP46 models. In both tumor models, the DNA topology analysis reveals tight contacts forming an anti- diagonal pattern on the contact map at the *PRAME* locus (Figure 6C). Such tight contacts could be caused by a hairpin structure of a single DNA loop.

Analyzing the ENCODE resource of transcription factor ChIP-seq experiments [53], [54], 128 transcription factor recognition sites were identified between *ZNF280A/B* and *PRAME* corresponding to 85 different DNA binding factors involved in chromatin and transcription regulation. Among these, 20 are also significantly upregulated in our gene expression analysis (Supplementary Figure 12). Expressed and upregulated *PRAME* DNA binding factors include cohesin components RAD21 and SMC3, which may contribute to this hairpin like topology. Chromatin organization modifiers such as DNA binding helicases CHD1 and CHD2, were also found upregulated in our analysis. All of these proteins are associated with chromatin remodeling and may contribute to *PRAME* overexpression in our UM models.

Overall, we have identified a novel pattern of chromatin conformation associated with striking overexpression of PRAME in UM.

## Discussion

We report an extensive multi-omics approach comparing two aggressive UM models with short-term cultured normal choroidal melanocytes. The multi-omics analysis includes whole genome somatic mutations, transcriptome, copy number, methylome, DNA optical mapping, FISH, histone modifications, and DNA topology analysis by Hi-C. The MP41 tumor does not display any identifiable BSE event, expresses BAP1 and is classified in the TCGA copy number group enriched in SF3B1-mut UM. The second tumor sample, MP46, belongs to the TCGA high-risk group and does not express BAP1 despite the absence of mutations in the BAP1 coding sequence.

Although disomy 3 UM patients display a favorable outcome some of them develop metastases. This is the case of the patient from whom MP41 was derived. Interestingly in the TCGA cohort a third of disomy 3 UM samples (13 out of 38) do not display a BSE event whereas the absence of BSE event is rare in the monosomy samples (3 out of 42). Given the absence of a BSE mutational event in MP41 we carefully analyzed the whole genome sequencing data for mutations potentially explaining tumor progression in MP41. In addition to *GNA11* mutation, a nonsense truncating mutation in *KMT2C* was predicted as an oncogenic driver. This gene is altered in about 5% of cancers but *KMT2C* mutations have not been described in UM [55]. The RNA level of *KMT2C* in MP41 is not significantly different of that displayed by NM and we observed a very high expression of this gene in MP46 (data not shown). Further functional studies restoring the wild-type allele of *KMT2C* in cell lines derived from MP41 need to be performed to explore the potential contribution of the nonsense *KMT2C* mutation in UM. MP41 may thus serve as an informative model of aggressive UM outside the classical BSE context.

Our gene expression profiling compared to normal uveal melanocytes revealed several interesting patterns of gene expression changes. The list of differentially expressed genes can help to identify new players in UM oncogenesis and potential new therapeutic targets. Interestingly, *PLCB4* and *RASGRP3* were among the top 50 upregulated genes in both tumor models. Whereas activating mutations of *PLCB4* have been described in UM, our data suggests that high expression can also contribute to over-activation of the PKC pathway. The increase in expression of RASGRP3 in UM samples is also of particular interest given that this protein has been shown to mediate MAPK pathway activation in UM [8], [9]. In the current study we have identified an additional GPCR downstream pathway gene, the Rap guanine nucleotide exchange factor *RAPGEF4* which is significantly upregulated in both UM PDX as compared to NM. Further functional studies must be conducted to evaluate the potential role of this protein in UM.

UM is classically considered to display a relatively simple pattern of karyotypic alterations in comparison with other solid tumors. Interestingly DNA repair is among the most deregulated pathways in our dataset. The TCGA consortium described an activation of DNA damage repair in transcription-based cluster 4, which is enriched for patients with the poorest prognosis. Consistent with activation of these pathways at the gene expression level, our structural DNA analysis combining optical mapping and FISH revealed multiple chromosome aberrations including intra and inter chromosomal translocations, insertion-duplications, telomeres shortening and telomere aberrations. Importantly, we observed chromosomal aberrations by optical mapping and FISH approaches in six other UM cell lines strongly suggesting that genomic / chromosomal instability is a hallmark of aggressive UM.

Bi-allelic inactivation of *BAP1* is associated with an increased risk of metastasis in patients with uveal melanoma. In this work, we describe a novel genomic source of *BAP1* deficiency consisting of a deletion on the *BAP1* promoter and boundary hyper methylation. A targeted NGS approach identified two other cases of UM displaying deletion in the same genomic area resulting in the lack of *BAP1* expression indicating recurrence of this mechanism. Given the correlation of *BAP1* deficiency with the risk of developing metastases, our results indicate that in UM patients it will be informative to look at the *BAP1* promoter when mutations or indels in the coding sequence are not detected. This has been recently implemented in Institut Curie where NGS is performed on UMs as part of the National initiative “*France Medecine Genomique 2025*”. Within this initiative, we have recently detected an additional UM case displaying the deletion of the *BAP1* promoter.

DNA topology was investigated by using an in situ Hi-C approach. Most changes in gene expression between NM and our two UM models were not accompanied with changes in compartment status. An important exception to this general pattern could be observed at the PRAME locus, where striking over-expression is accompanied by compartment switching and a qualitative change in the intra-TAD contact pattern with an anti-diagonal pattern of associations suggesting a potential hairpin-like structure associated with gene activation. This finding highlights the relevance of using a multi-omics approach that combines chromatin confirmation alterations and RNA expression. Further investigation of repositioned compartments identified in our comparison could help address questions of compartment specificity recently identified in other tumor types [56].

In summary, our study illustrates how multi-omics integrative approaches conducted in a limited number of samples can improve the understanding of tumorigenesis and reveals two novel mechanisms of gene expression dysregulation in UM with potential clinical implications.

## Methods

### Isolation of UM cells from PDX

As described before [19], [20], [22], MP41 and MP46 xenograft tumors were harvested before they reached a volume of 1 cm3 following ethical rules, and processed immediately for dissociation, immunolabelling and sorting based on Petit et al protocol[57]. To avoid isolation of heterogeneous UM cell populations in batches of experiments, immunostaining was conducted with the anti Muc18 containing a rabbit Fc: clone 8H2rFc, also with anti CEACAM1 (8G5hFc) and anti NG2 (14A7hFc) as characterized previously [22]. Secondary antibodies were anti rabbit FC-AF647nm to reveal Muc18 labelling, and anti-human Fc to reveal at the same time CEACAM1 and NG2 labelling. Cell sortings were conducted with the help of cytometry platforms of Institut Curie on single live cells. In total, 15-20 mice were grafted per model, generating 225x10^6^ cells for MP41 and 133x10^6^ cells for MP46. Twelves batches of dissociation and cell sorting were carried out for MP41 and 10 batches for MP46 generating in total 225x10^6^ and 133x10^6^ cells respectively for MP41 and MP46. After each cell sorting, a fraction of cells was kept for DNA and RNA extraction, and most cells were fixed according Rao et al protocol.

Normal uveal melanocytes from two healthy donors and 3 preparations obtained after enucleations were dissociated and maintained in culture from choroidal membranes from Pr Simon Saule’s lab and Dr Geraldine Liot. After enzymatic (with collagenase) and mechanical dissociation, primary cells were cultured in HAM/F12 supplemented with 10% FBS, Penicillin 100 U/mL, Streptomycin 100 U/mL, 2mM L-glutamine, 2.5µg/mL Amphotericin B. Extemporaneously completed media was supplemented with 0.1mM IBMX, 10ng/µL of Cholera toxin (10 ng/ml final), and 10mg/mL βFGF and filter with a 0.22µm filter. Cell were maintained in a humidified atmosphere with 5% CO2 at 37°C, and culture medium was exchanged twice a week.

### Quality control of isolated cells

To monitor each fraction of MP41/MP46 isolated cells, DNA and RNA were extracted as described before, and tested for chromosomal copy number alterations and gene expression using Affymetrix microarrays respect to previous analysis done on primary tumor or pdx samples [22].

Based on this analysis, pools of MP41 cells and MP46 cells were used for whole genome sequencing, RNAseq, DNA methylation, *in situ* Hi-C, ChIPSeq experiments, and to allow replicates of in situ Hi-C and ChIPSeq analyses.

### Whole genome analysis

Genomic DNA was extracted using QIAamp DNA Mini kit, and quality controls were achieved using a Nanodrop ND1000 to evaluate DNA purity and Qubit™ dsDNA BR/HS Assays to evaluate double strand DNA concentration. 250ng of gDNA from MP41 and MP46 sorted cells were characterized using Affymetrix/Thermo Cytoscan HD microarrays to monitor copy number and LOH. Next, two micrograms of MP41 and MP46 DNAs were used to prepare paired-end 100bp Illumina libraries for whole genome sequencing. Genomic DNA from healthy surrounding tissue was sequenced according to approval by ethic committee of Institut Curie to filtered germline mutations.

Whole genome sequencing was conducted on genomic DNA extracted from PDX derived UM cells and from normal tissue preserved from enucleation. Illumina short read sequencing was achieved in two-separated runs due to availability of DNAs. The alignment of the sequenced reads to hg19 was performed with Burrows-Wheeler Aligner (BWA-MEM v0.7.10). MuTect2 from the Genome Analysis Toolkit (GATK v3.5) was used in "Tumor with matched normal" mode to call somatic variants. Somatic point mutation analysis of whole genome sequencing revealed 1348 and 1186 single nucleotide variants in MP41 and MP46 respectively representing less than one somatic mutation per Mb (0.42 and 0.37 SNV/Mb) as observed in UM [18]. Detected variants with a frequency in the normal sample greater than 20% were filtered out. Variants in UM models were selected according coverage (>10), read counts (>20) and allele frequency (>20%) in UM models. Cancer Genome Interpreter (CGI, [23]) and VarSome tools (containing 10 pathogenic predictions, [24]) were combined to characterize variants.

**<u>Whole genome DNA methylation analysis</u>** was carried out with a Cambridge Epigenetics kit (TrueMethyl kit) that corresponds to an oxidative bisulfite reaction, to identify and analyze only 5-methylcytosine (5-mC). Briefly 400ng of genomic DNA were used to first perform an oxidation reaction, that converts 5- hydroxymethylcytosine (5-hmC) to 5- formylcytosine(5fC), thus after a bisulfite conversion, unmodified C and 5FC will be converted into uracils and sequenced as thymines contrary to 5mC that remains sequence as a cytosine. Libraries were sequenced on an Illumina Hiseq as paired end 100bp. Paired-end reads were trimmed with fastx toolkit v0.0.13 with these parameters : -f 8 -Q 33. Adapters were removed using Cutadapt v1.8-2. Cleaned reads were aligned with bismark v0.12.5 with default parameters onto the Human reference Genome Hg19. Only reads mapping uniquely on the genome were conserved. Methylation calls were extracted after duplicate removal. Only CG dinucleotides covered by a minimum of 5 reads were conserved for the rest of the analysis. Differentially methylated regions (DMR) were assigned when a methylation difference of 30% occurs at least on 10 CpGs in a minimum of 500bp windows; windows are merged if distant between two DMRs is less than 500bp. DMR calling was performed using the Bioconductor package DSS [58].

### Chromatin structure analysis

We performed *in situ* Hi-C in duplicate for 3 normal melanocytes (NM) and 2 UM models (MP41 and MP46) as described by Rao et al [59] Briefly, after cell sorting of MP41 and MP46, 3x10^6^ NM and UM cells were fixed in 1% formaldehyde and stored at -80°C until further processing. Fixed cells were permeabilized, and an overnight digestion with MboI was conducted. DNA overhangs were filled in in the presence of biotin-14- dATP before proximal ends were ligated with T4 DNA ligase for 4 hours. After crosslink reversal with Proteinase K, DNA was purified using phenol/chloroform, quantified and sheared to a size of 400-500bp. Next, biotinylated DNA was pulled down with Dynabeads MyOne streptavidin T1 beads (Thermo Fisher) and DNA was repaired for 30 minutes with a mixture of T4 DNA ligase (NEB), T4 DNA polymerase I (NEB), large fragment of DNA polymerase I (NEB), and T4 Polynucleotide Kinase (NEB). Beads were washed and separated on a magnet before dA-tailing with Klenow exo minus (NEB). A final wash was performed before Illumina adaptor ligation. PCR amplification with Illumina primers was performed for 12 cycles and product was collected with AMpure. An equimolar pool of libraries was sequenced on an Ilumina HiSeq - rapid Run, generating 250-436x10^6^ read pairs. Hi-C data were processed using HiC-Pro [60] before comparing 3 different normalization algorithms (CAIC, LOIC and ICE methods, [43]).

### DNA Optical mapping and cytogenetics analysis

SVs analysis was conducted with Bionano DNA optical mapping from 1.5x10^6^ MP41 and MP46 cell pellets. A direct labelling on CTTAAG motif (DLE1) was conducted according to Bionano recommendations. Labelled DNA were analyzed on a Saphyr system. A De novo assembly was carried out using the Bionano serve 1.6. Molecule N50 was 407.8kbp for MP41, label density was 16.3 per 100kbp and effective coverage of the assembly was 71.9X. For MP46, molecule N50 was 325.6kbp, label density was 16.9/100kbp and effective coverage of assembly was 84.5X.

Telomere and centromere staining followed by M-FISH technique were applied on cytogenetics slides after colcemid (0.1µg/mL) treatment of MP41 and MP46 cells as described previously [34], [61] to identify numerical and structural chromosomal alterations as well as telomere instability. Briefly, UM cells were cultured in T75 in DMEM with 10-20% SVF depending on models (10% SVF: MP41, Mel202, OMM1, OMM2.3; 20%: MP46). Forty-eight hours after passage, medium was supplemented with colcemid (0.1 µg/mL) for a 3h incubation to arrest dividing cells in metaphase. Cells were harvested, washed, suspended in 10mL DMEM with 0.075 M potassium chloride (Merck, Kenilworth, NJ, USA) and incubated for 20 min in a 37°C water bath (hypotonic shock) and fixed as previously described [34]. Next a telomeres and centromeres followed by M-FISH technique (Metasystems probes, Germany) were performed as previously described [62]. The quantification of telomere FISH signal intensity in interphase cells was performed using Metacyte software (MetaSystems, version 3.9.1, Altlussheim, Germany) and TeloScore Software (Cell Environment, Evry, France).

The mean fluorescence intensity (FI) of telomeres was automatically quantified in 10,000 nuclei on each slide. The settings for exposure and gain remained constant between captures. The experiments were performed in triplicate.

Analysis of metaphase spreads allowed detection of telomere abnormalities and chromosomal aberrations using ChromoScore Software (Cell Environment, Evry France) and Isis software (MetaSystems, Altlussheim, Germany). The images of metaphases were captured using automated acquisition module Autocapt software (MetaSystems, version 3.9.1) and a ZEISS Plan-Apochromat 63x/1.40 oil (Zeiss, Oberkochen, Germany) and CoolCube 1 Digital High Resolution CCD Camera (MetaSystems, Altlussheim, Germany) with constant settings for exposure and gain.

For each UM model, telomere and chromosomal aberrations were analyzed automatically on 100 metaphases. The scored telomere abnormalities were (i) sister telomere loss, likely occurring in G2, and defined as a telomere signal-free end at a single chromatid, (ii) telomere deletion defined as the loss of two telomere signals on the same chromosome arm (likely resulting from the loss of one telomere in G1/S), an aberration considered to represent double strand breaks, leading to activation of DNA damage response. The scored chromosomal aberrations were dicentric chromosomes, centric rings, translocations, insertions and deletions.

### Whole transcriptome analysis

Total RNA was extracted using miRNeasy kits following supplier recommendations, including a DNAse step. Quality controls were achieved using a Nanodrop ND1000 to evaluate RNA purity and concentration, and on automated electrophoresis to monitor RNA integrity (Bioanalyzer RNA 6000 Nano/Pico kits). PolyA RNASeq was conducted on total RNA (RIN>7), treated with DNAse. An absolute fold change higher than 1.5 and a p-value below to 0.05 were selected as parameters for detecting differentially expressed genes. Splicing analysis was conducted with 5 different pipelines: deFuse, SOAPfuse, JAFFA, FusionCatcher, TopHat-Fusion. Fusion RNAs were identified present in at least 2 algorithms, and found in at least 2 replicates per model.

### Histone Modifications

ChIPSeq against H2AUb, H3K4me and H3K27me3 were conducted in simplicate in NM, MP41 and MP46 as published in [52]. ChIpSeq against H3K27Ac and CTCF, were conducted in duplicated in MP41 and MP46 to implement multiomics analysis. 5 million cells were fixed according the protocol used for in situ Hi-C experiments for H2AUb, H3K4me3, and H3K27me3, and for H3K27ac and CTCF. The chromatin was prepared using the iDeal ChIP seq kit from Diagenode for Transcription Factor protocol. Shearing conditions were setup as 10 minutes using the following settings: 10 cycles of 30’’ [ON] 30’’ [OFF]. The shearing efficiency was monitored after reversion of the crosslinking and purification of the DNA. For increased sensitivity, an automated capillary electrophoresis system Fragment Analyser was used for chromatin shearing assessment (High sensitivity NGS fragment kit). ChIP assays were performed as defined in the optimizations using 10 μg or 1μg of chromatin per IP with the optimal antibody quantity resulting in the higher enrichment and lower background (CTCF 1μg, H3K27ac 1μg) IPs with a negative control isotype (IgG) were performed in parallel. For each sample, a library preparation was performed on ChIP and input DNA using the MicroPLEX v3 protocol. A control library was processed in parallel with the samples using a control Diagenode ChIP DNA. Five cycles of pre-amplification were performed and 1 μl of each library was analyzed by qPCR in order to determine the optimal amplification cycles required to obtain enough DNA for sequencing. Libraries were then loaded on Fragment Analyzer to check if enough material was generated. After the amplification, the libraries have been purified using AMPure beads and eluted in Tris. Then, the purified libraries were quantified using the Qubit ds DNA HS kit and analyzed on the Fragment Analyzer to assess their size. Using the quantification values from the Qubit and the size measurement generated by the Fragment Analyzer, the molar concentration of each library was calculated.

The quality control of the fastq files was performed using FastQC. The sequences were aligned on hg19 assembly using bowtie2. Duplicates were marked and filtered out using Picard tools MarkDuplicates and samtools. The peak calling was performed using macs2 callpeak function. The parameter --broad was used for the Histone samples, not for the transcription factor samples. The affinity binding scores were obtained using DiffBind package in R, TMM normalization was applied. Peaks found in at least 50% of the samples from the same condition were kept. The peaks were annotated using FAST DB.

## Acknowledgements

The authors thanks the members of the Institut Curie animal facility involved in UM projects, the members of the cytometry platform of Institut Curie and Hospital Saint Louis in Paris for their support of UM cell sorting, the members of the recombinant protein and nanobodies platforms of Institut Curie, the Pathex platform of Institut Curie for BAP1 immunochemistry, the biological resources center for providing UM samples for the BAP1 analysis. The author thank the Unit of Somatic Genetic at the hospital of Institut Curie for the analysis done on NM. The authors thank the genomics platform of Institut Curie for assistance in STR profiling, copy number analysis, DNA optical mapping of UM samples (Audrey Rapinat, Matéo Bazire), with the support of the Region Ile France (SESAME 2019 grant). High-throughput sequencing was performed by the ICGex NGS platform of the Institut Curie supported by the grants ANR-10-EQPX-03 (Equipex) and ANR-10-INBS-09-08 (France Génomique Consortium) from the Agence Nationale de la Recherche ("Investissements d’Avenir" program), by the ITMO-Cancer Aviesan (Plan Cancer III), and by the SiRIC-Curie program (SiRIC grant INCa-DGOS-4654). This work was supported too with a grant from the National Human Genome Research Institute to Job Dekker (HG003431). Job Dekker is an investigator of the Howard Hughes Medical Institute. The authors also thank the Bionano Genomics company and the GENTYANE platform in Clermont Ferrand for the first analysis of MP41 and MP46 models using optical mapping.

## Author Contributions

DG, EH, SRR, JJW, conceived and designed the study. DG, NS, CR, LVG, MB, TC, SB, RM, EJ, AN, collected the data. DG, ESA, NS, AJ, PDLG, GL, SS, AT, DB, RM, EJ, AN, MHS, JHG, JD, JW, JMP, LVG, SRR and JJW analyzed and interpreted the data. DG, SRR and JJW wrote the manuscript. All authors reviewed and approved the final version of the submitted report.

## Data availability

Supplementary table 1 is available upon request. Sequencing data generated in this work will be deposited on a public portal.

---

Supplementary Figure 1: Localization of shared regulated genes in MP41 and MP46 vs NM. Localization of the 2334 upregulated genes and the 3066 down regulated genes are represented respectively in red and green on a chromosome view.

Supplementary Figure 2: Main regulated pathways from the regulated genes shared in MP41 and MP46 vs NM. Gene names are colored depending on their regulation: non-regulated gene are in white boxes, upregulated genes in red boxes and downregulated genes in green boxes. Light colors indicate a significant regulation in MP41 vs NM or in MP46 vs NM, and dark colors indicate a significant regulation observed in both comparisons. A) Cell cycle pathway (HSA04110, KEGG). B) Homologous recombination (HR) pathway (HSA03440, Kegg). C) Fanconi Anemia pathway (FA, HSA03460, Kegg). D) Non-homologous end joining pathway (NHEJ, HAS03450, Kegg). E) P53 signaling pathway (HSA04115, KEGG). **F.** Apoptosis pathway (HSA04115, KEGG).

Supplementary Figure 3: Telomere analysis of MP41, MP46 and normal fibroblasts. A) Telomere fluorescence intensities of the 3 models is function of cell area. B) Rate of telomere loss per chromosome in MP41, MP46 compared to controls. Yellow stars highlight telomere loss on 3 chromosomes of MP41. C) Frequency of telomere loss and deletion per chromosome in MP41 and MP46. D) Example of telomere defects as interstitial telomere observed on dicentric (6;8;17) in MP41, and on a derivative chr15. Telomere deletion on dic(14;16), i8q, and chr6, and chr 14q are illustrated. E) A dic(1)t(1;11;6;8) and a ring chromosome probably coming from a breakpoint on 1q21 leading to a ring chromosome (r(dic(1;11;6;8)). F) A ring chromosome in MP46 (r(21)).

Supplementary Figure 4: FISH analysis of chromosome 6 and 8 in MP41 model. A) Specific probes (Empire Genomic, USA) designed for the detection of 4 genes: *NCOA7* (orange), *RSPO3* (blue), *GPAT4* (green), and *POMK* (red). FISH probes are represented on the optical map showing the t(6;8) generated from the Bionano analysis. B) the presence of *RSPO3* and *NCOA7* in chromosome 6 without rearrangements (C) dic(6;8;17) showing the presence of *GPAT4* and *POMK* genes in addition to *RSPO3* and NCOA7 in the breakpoint of this aberration. (D) dic(6;8;16) showing the presence of *GPAT4* and *POMK* genes in addition to *RSPO3* and *NCOA7* (E) the i(8q) shows the presence of *GPAT4* and *NCOA7*. (F) dic(1;11;6;8) shows the insertion of the *GPAT4* and *NCOA7* in the breakpoint of this rearrangement (G) possible mechanisms in the formation of these rearrangements implicated chromosome 6 and 8.

Supplementary Figure 5: DNA optical mapping and multispectral FISH highlight chromosome aberrations and DNA rearrangements in UM models. A) Log2 ratio generated from Cytoscan HD microarrays of chr8 in UM models. B. Circos plot generated from DNA optical mapping illustrated main SVs as translocations. C. Telomere aberrations per chromosome on UM cell models as MP41, MP46 Mel202, MM66, OMM1, OMM2.3 and normal fibroblasts as controls. Main derivative and dicentric chromosomes of D) Mel202, E) MM66, F) OMM1, G) OMM2.3. FISH and TC pictures of derivative and dicentric chromosomes are presented. Blue triangles point small parts of a donor chromosome.

Supplementary Figure 6: Identification of *BAP1* deletion leading to *BAP1* loss of expression. Gene expression detection based on RNAseq analysis (FPKM counts) of A) *BAP1* and B) *PHF7* in NM, MP41 and MP46 samples. C) Targeted sequencing based on tilling amplicons (250bp) to characterize *BAP1* mutations in a series of 53 UM recently grafted for new UM PDX at Institut Curie. D) Detection of *BAP1/PHF7* promoter deletion in IGV. Blue track corresponds to NM sample, yellow track corresponds to MP41 and red track corresponds to MP46. Gray tracks correspond to new UM cases with a low coverage on *BAP1* promoter associated to a loss of BAP1 protein expression assessed by IHC (E).

Supplementary Figure 7: Example of *BAP1* promoter deletion identified in clinical daily practice. A) Identification of BAP1 promoter deletion (2.2kb) based on exome analysis. B. Validation of the loss of BAP1 expression in nucleus by IHC. C. Confirmation of the deletion of *BAP1* promoter by long range PCR in MP46 and new patient’s tumor.

Supplementary Figure 8: Comparison of WGS and Hi-C copy number profiles for MP41 and MP46. Two whole genome copy number views established with WGS (upper panel) or with Hi-C seq (lower panel) for A) MP41 and B) MP46. Copy number status as normal, gains and losses are shown respectively in green, red and blue.

Supplementary Figure 9: Effect on compartment interaction scores after different normalization methods applied to Hi-C data generated for MP41, MP46 and NM. Log2 Hi-C interaction scores are depicted across copy number status defined at a 250kb window resolution. Interactions score generated on A) raw data, or after B) ICE, C) CAIC and D) LOIC normalization. Per boxplot graph, each color corresponds to a specific copy number.

Supplementary Figure 10: Normalization of Hi-C Data and comparison of compartment interaction scores per type of compartment, and interaction score distribution per copy number status and number of A/B compartments. From each normalization methods of Hi-C data generated for MP41, MP46 and NM, interactions scores for all compartments were separated on upon A/B status, and undefined status. Log2 Hi-C interaction scores are depicted across copy number status defined at a 250kb window resolution. Interactions score from A) raw data, or after B) ICE, C) CAIC and D) LOIC normalization are compared for MP41, for MP46 and for NM. Per UM and NM panels, box plots and histograms show respectively compartment interaction scores and number of interactions. Green, blue and grey box plots and bars correspond respectively to euchromatin / A compartment, heterochromatin / B compartment, and not defined compartments.

Supplementary Figure 11: *PRAME* locus transcription factors (TF) upregulated in UM models. Hierarchical clustering of TF enriched at *PRAME* locus in ENCODE. Blue stars indicate significantly upregulated TFs in MP41 and in MP46 vs NM.

Supplementary Table 1: MP41 WGS analysis and list of SNV annotated

Supplementary Table 2: MP42 WGS analysis and list of SNV annotated

Supplementary Table 3: Somatic copy number alterations in UM models identified by WGS

Supplementary Table 4: Gene expression analysis between MP41 vs Normal Melanocytes (NM) and MP46 vs NM.

Supplementary Table 5: Number of genes in compartments. A: Euchromatin, B Heterochromatin; "AAA" letters correspond to A or B status in NM, MP41 and MP46 respectively.

