## Supplementary figures and tables for "Multi-omics comparison of malignant and normal uveal melanocytes reveals novel molecular features of uveal melanoma": Supp figures 220712.pdf

### Supplementary Figure 1: Localization of shared regulated genes in MP41 and MP46 vs NM

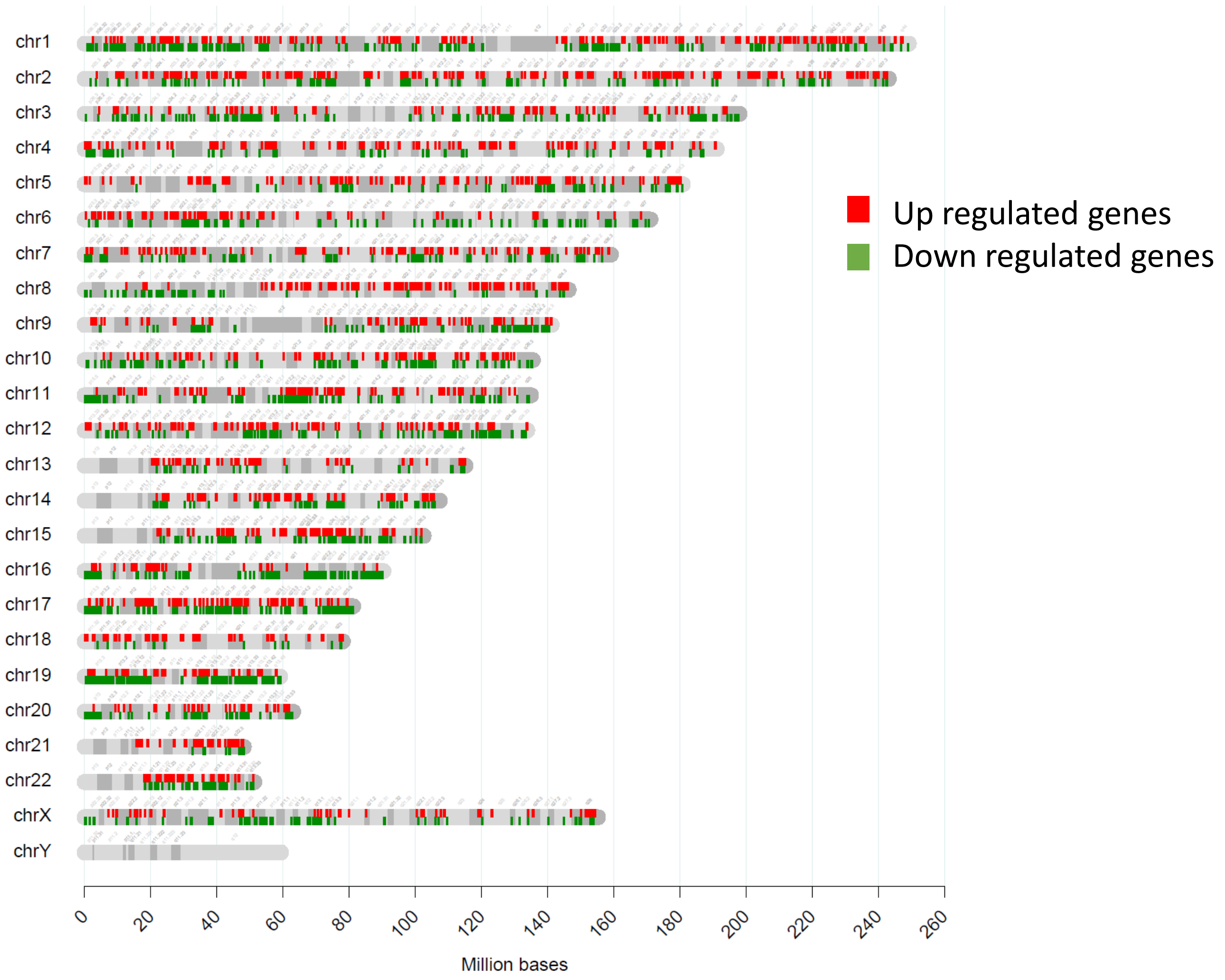

### Supplementary Figure 2: Main regulated pathways from the regulated genes shared in MP41 and MP46 vs NM

A. Cell cycle (HSA04110, KEGG)

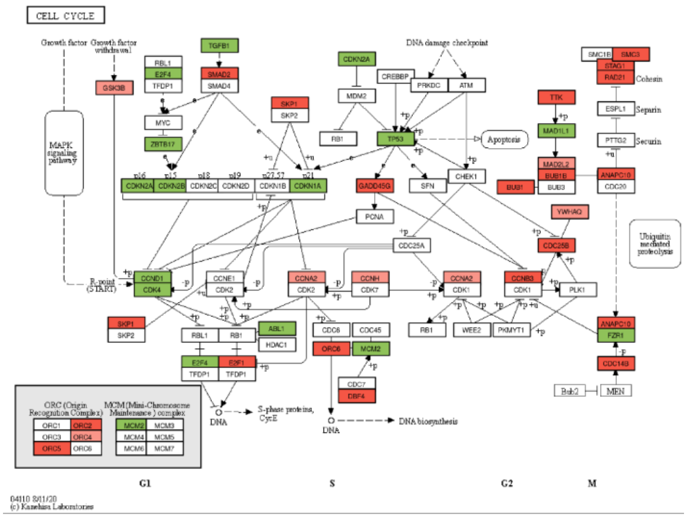

43 regulated genes out of 124 genes in the Cell cycle pathway, min p-value: $5.30 \times 10^{-07}$ (28 Up & 15 Down)

B. Homologous recombination (HSA03440, KEGG)

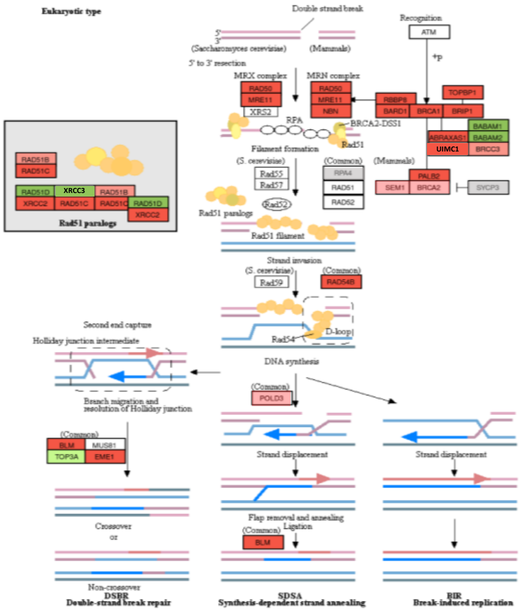

21 regulated genes out of 41 genes in the Homologous recombination pathway, min p-value: $5.99 \times 10^{-09}$ (17 Up & 4 Down)

C. Fanconi Anemia (HSA03460, KEGG)

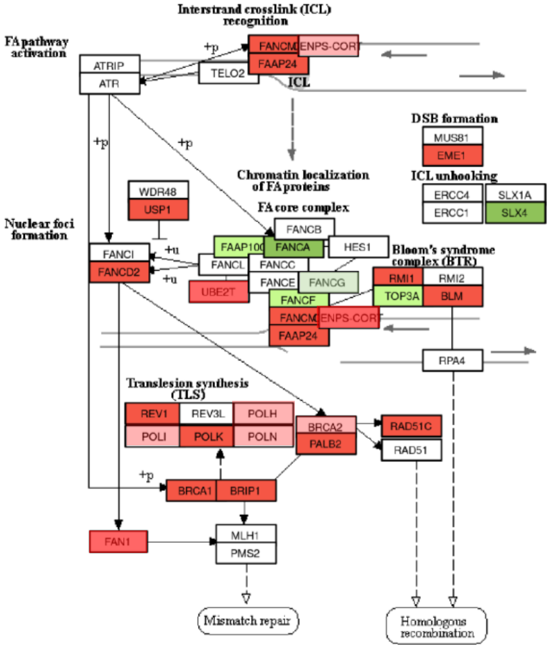


19 regulated genes out of 54 genes in the FA pathway, min p-value: $2.94 \times 10^{-4}$ (13 Up & 6 Down)

D. Non-Homologous End-Joining (HSA 03450, KEGG)

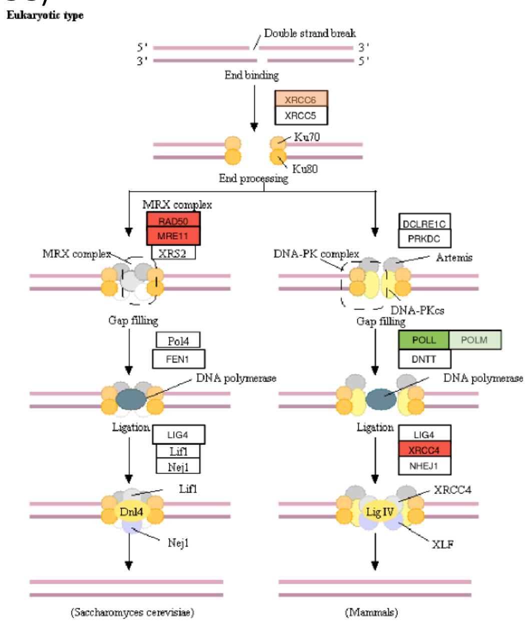


4 regulated genes out of 13 genes in the NHEJ pathway, min p-value: $8.22 \times 10^{-2}$ (3 Up & 1 Down)

### Supplementary Figure 2: Main regulated pathways from the regulated genes shared in MP41 and MP46 vs NM

**E. p53 signaling pathway (HSA04115, KEGG)**

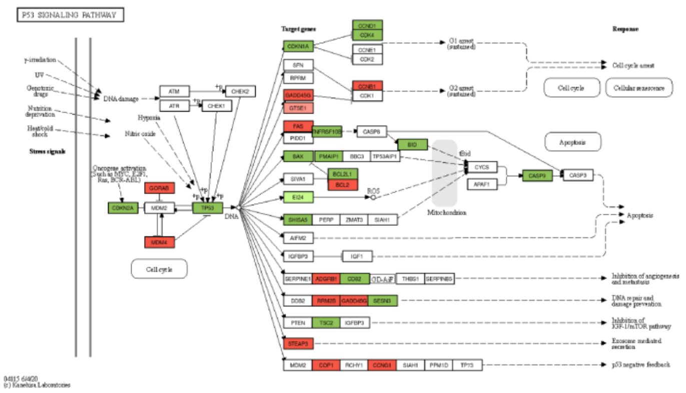

32 regulated genes out of 72 genes in the p53 signaling pathway, min p-value: $1.18 \times 10^{-6}$ (14 Up & 18 Down).

**F. Apoptosis pathway (HSA04210, KEGG)**

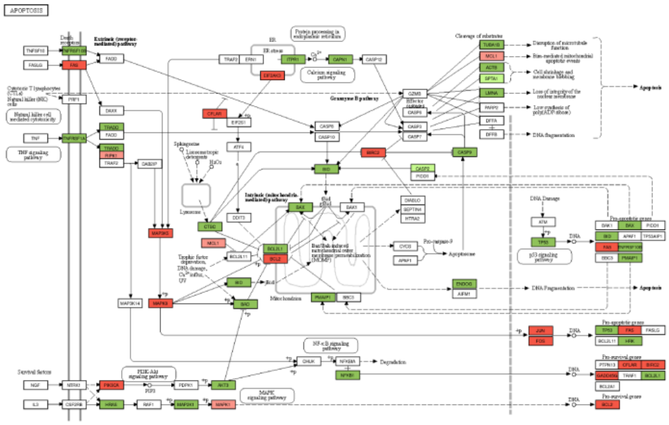

52 regulated genes out of 136 genes in the apoptosis pathway, min p-value: $2.47 \times 10^{-07}$ (20 Up & 32 Down)

Supplementary Figure 3: Evaluation of telomere instability based on T/C + M-FISH analysis.

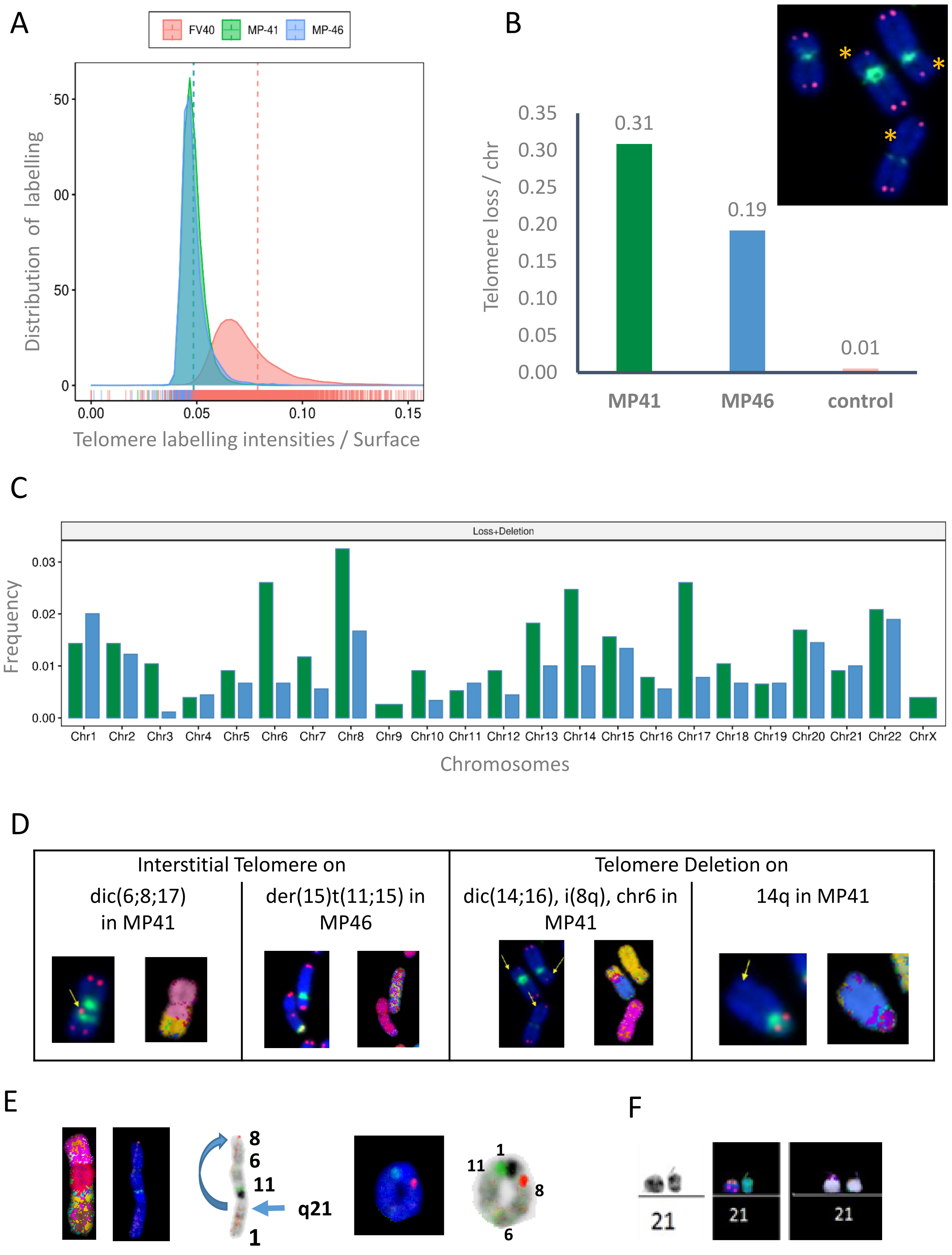

Supplementary Figure 4: FISH analysis of chromosome 6 and 8 in MP41 model.

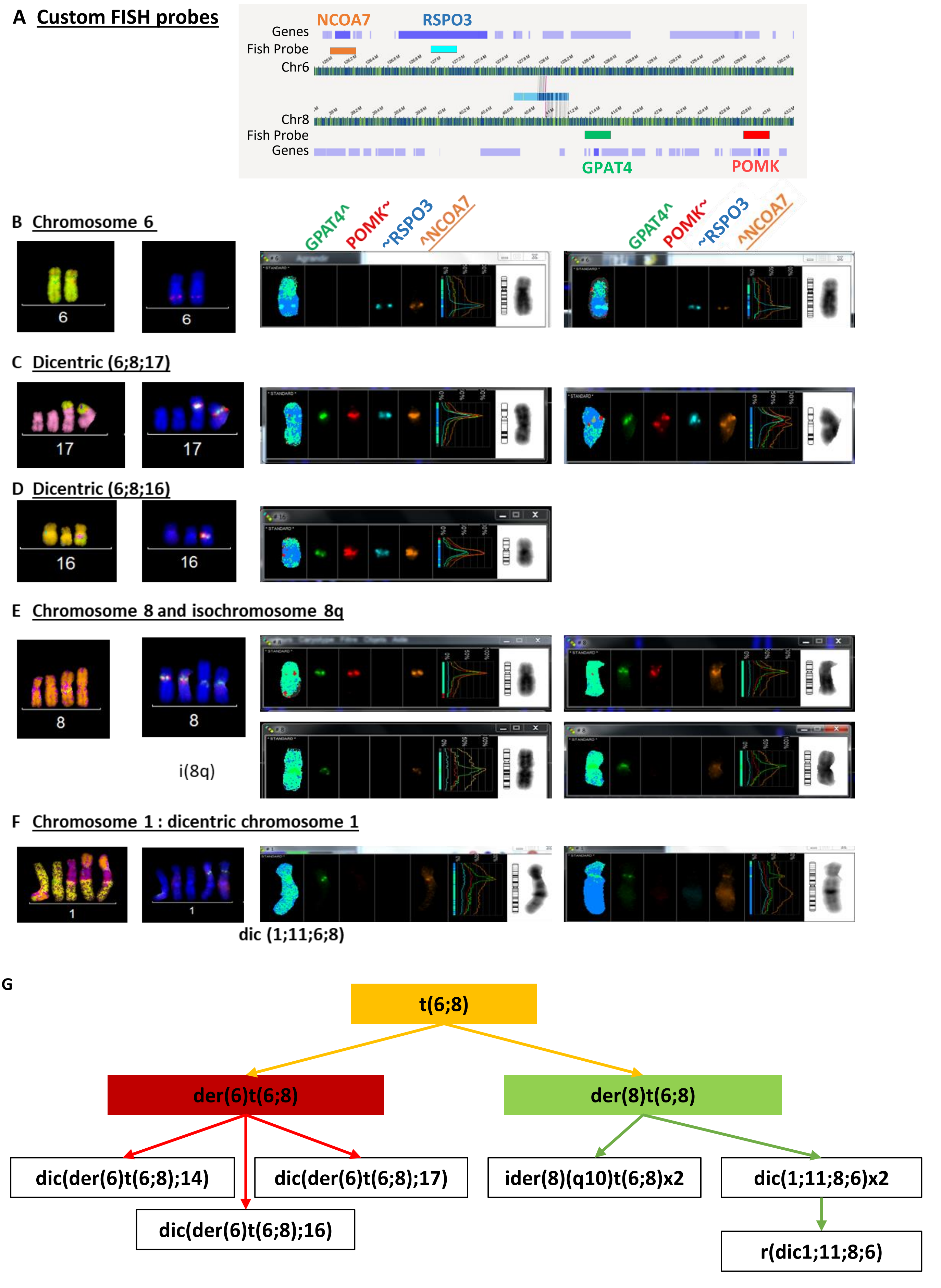

### Supplementary Figure 5: DNA optical mapping and multispectral FISH in UM models

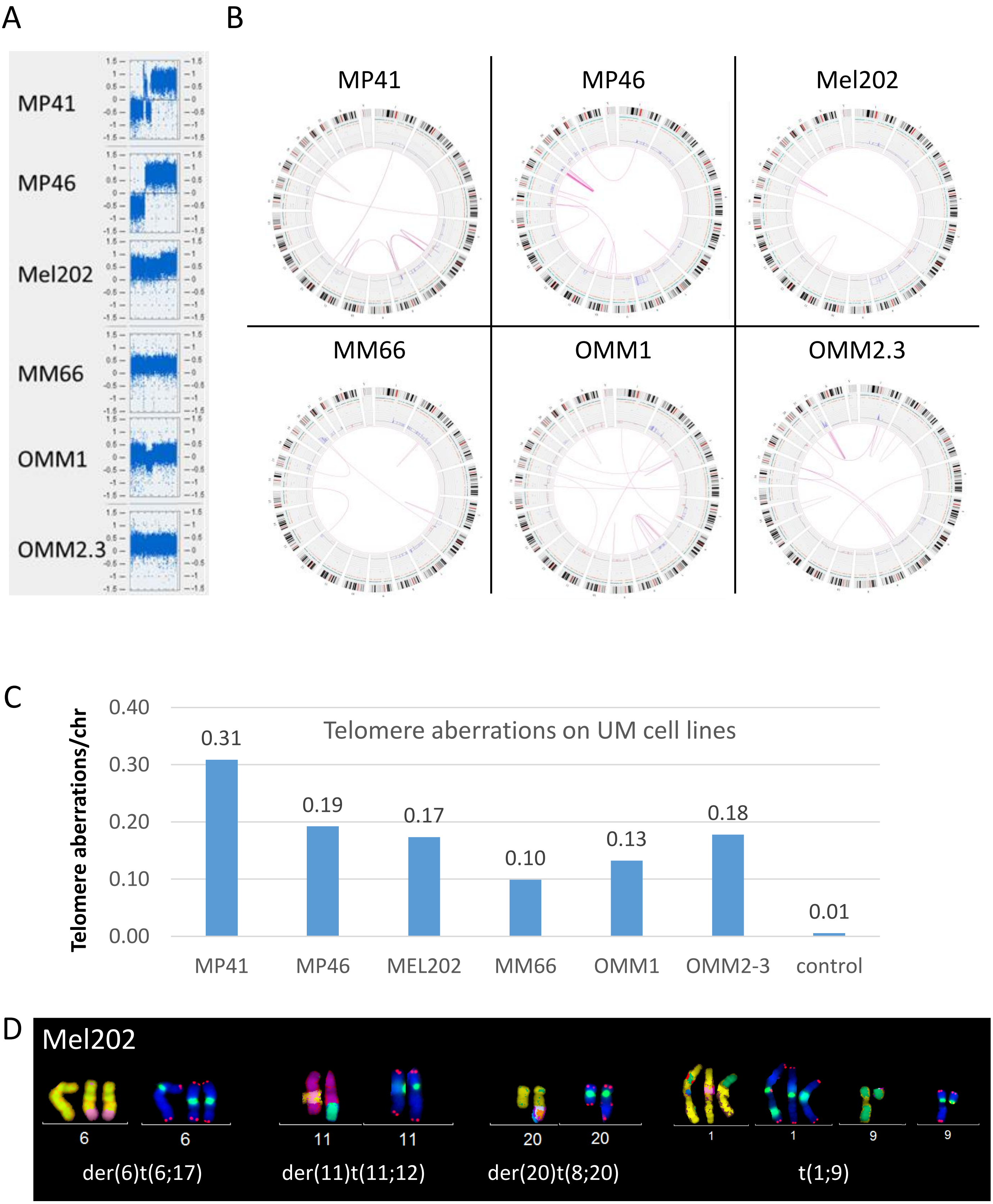

Supplementary figure 5: DNA optical mapping and multispectral FISH in UM models

E

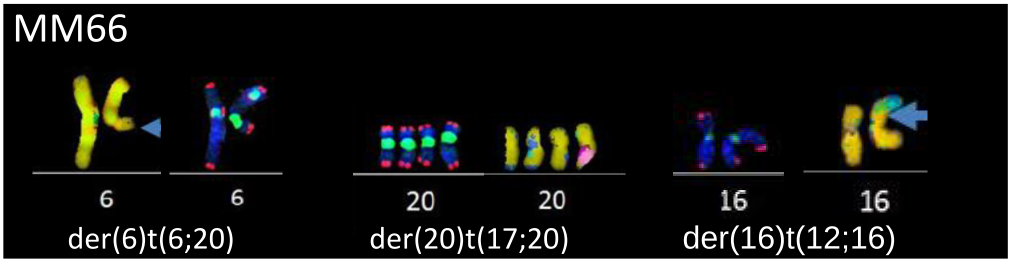


F

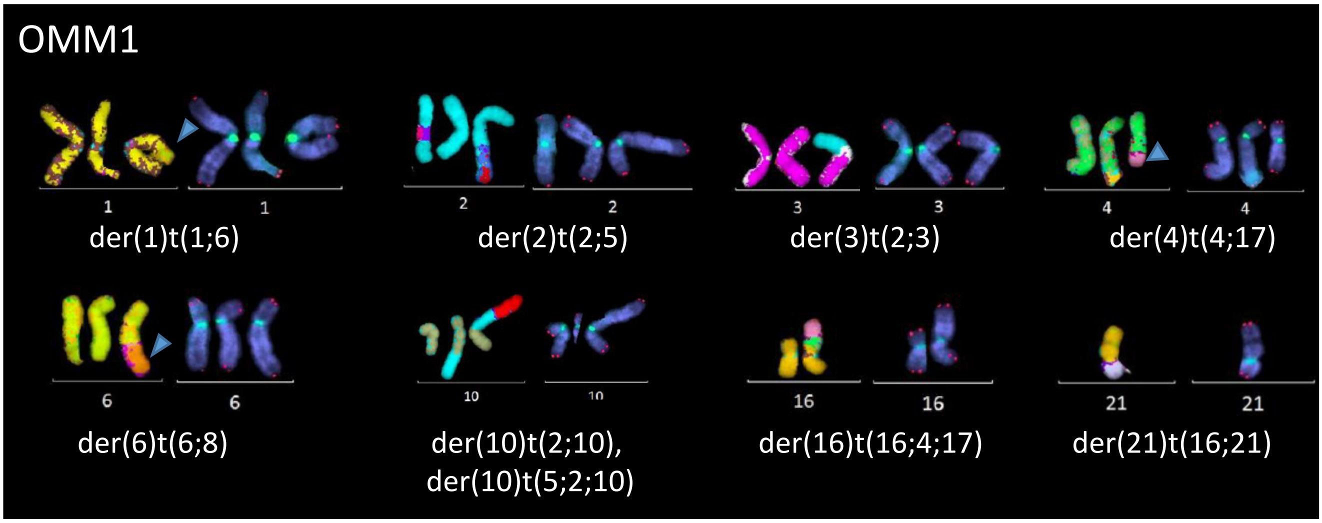


G

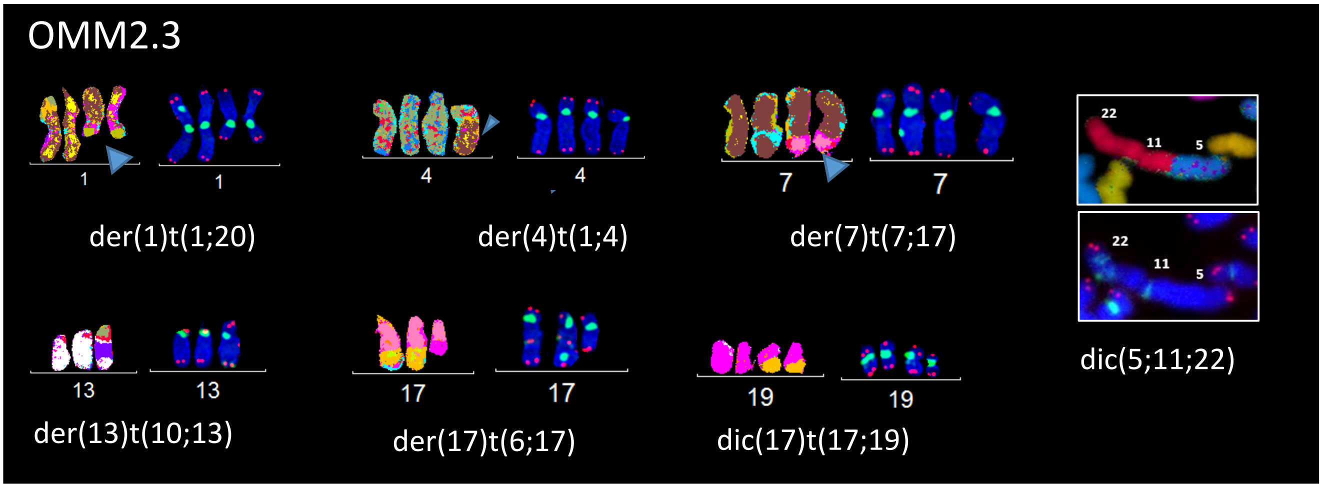

### Supplementary Figure 6: identification of *BAP1* deletion leading to BAP1 loss of expression

A. *BAP1* gene expression

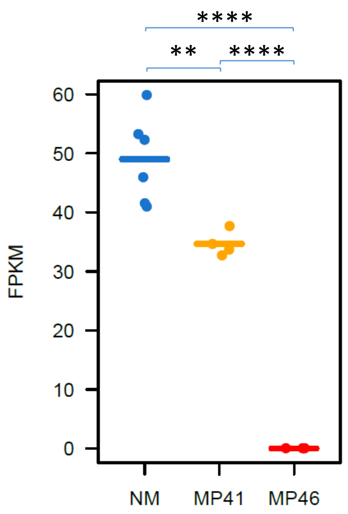


B. *PHF7* gene expression

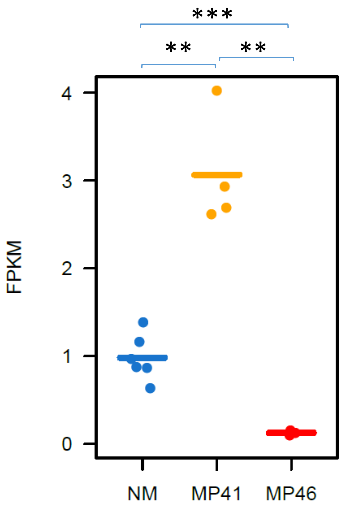


C. *BAP1* amplicon sequencing

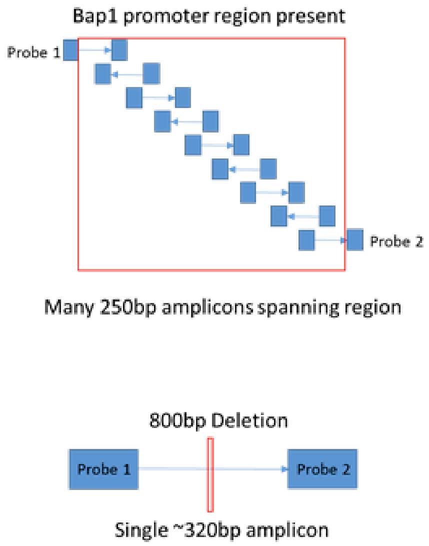


D. *BAP1* targeted DNA sequencing

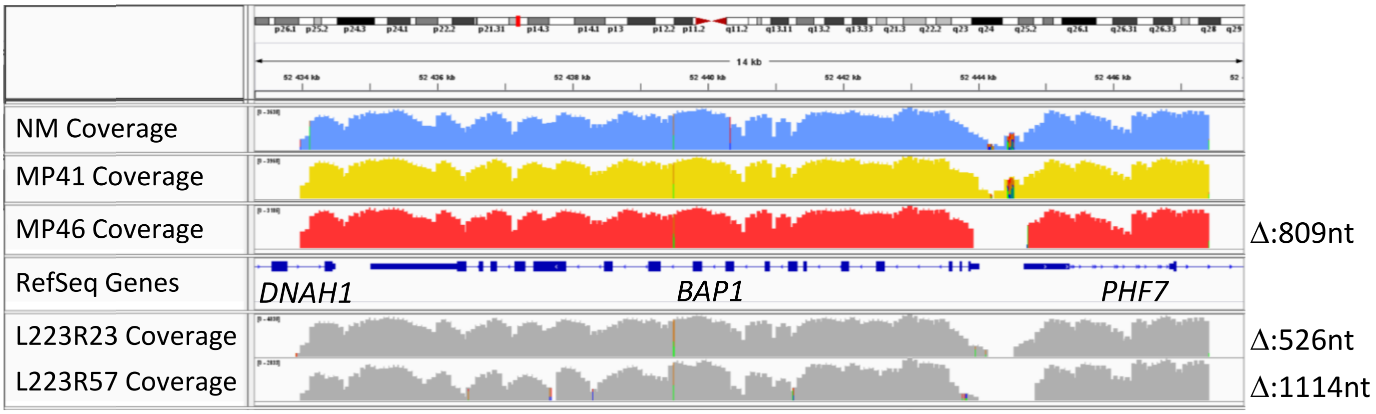


E. BAP1 IHC (40X)

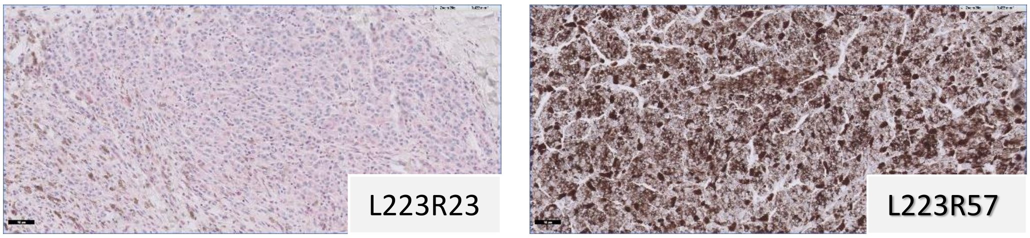

Supplementary Figure 7: Example of *BAP1* promoter deletion identified in clinical daily practice

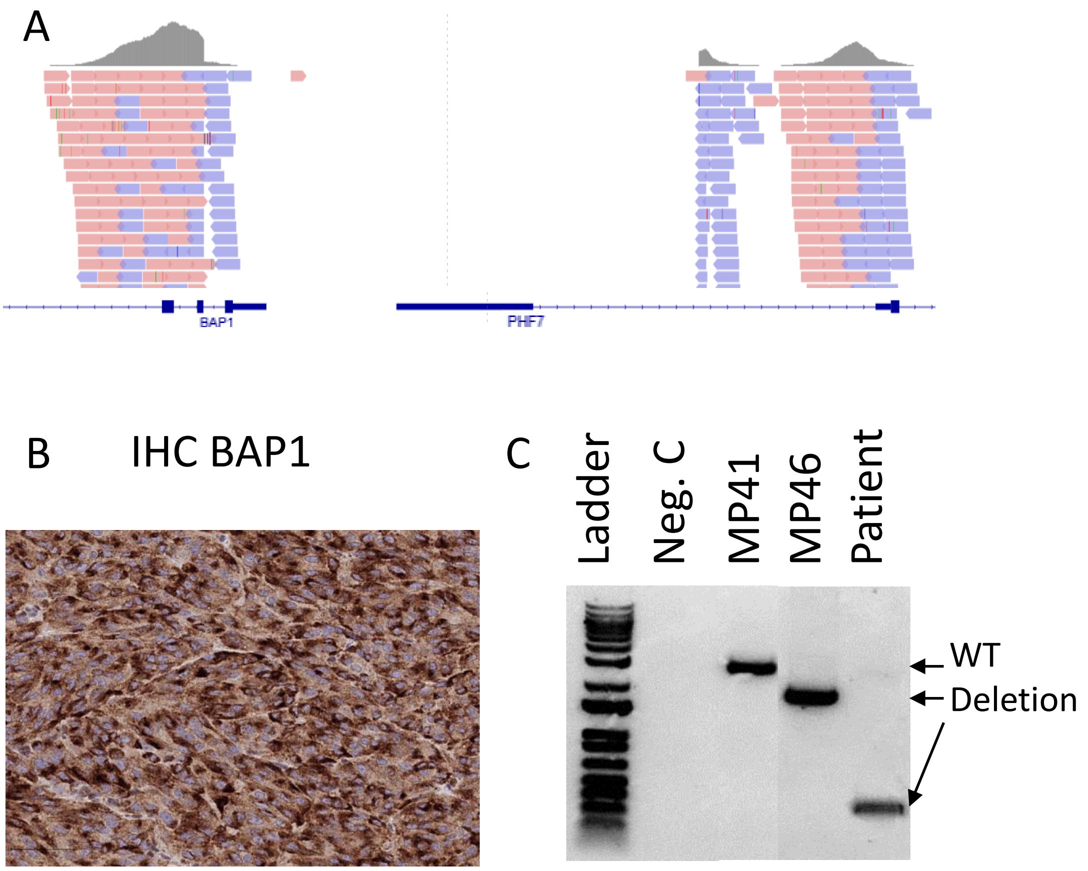

Supplementary Figure 8: Comparison of WGS and Hi-C copy number analysis for MP41 and MP46.

A

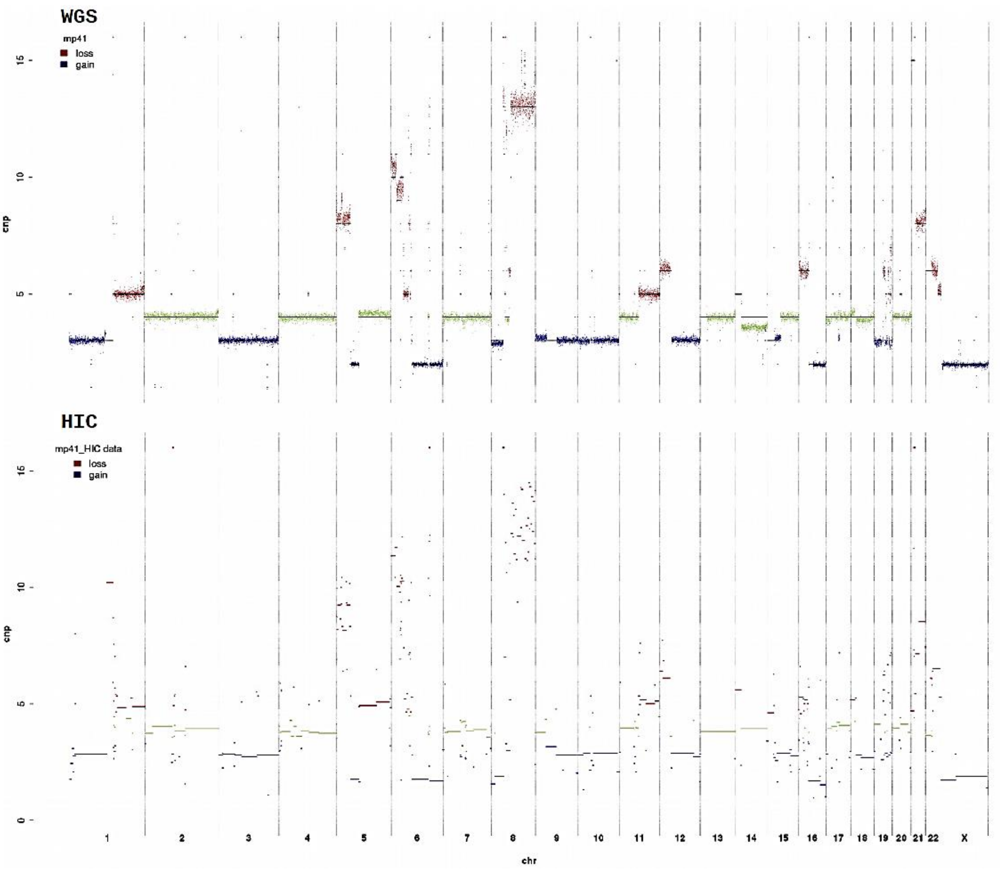


B

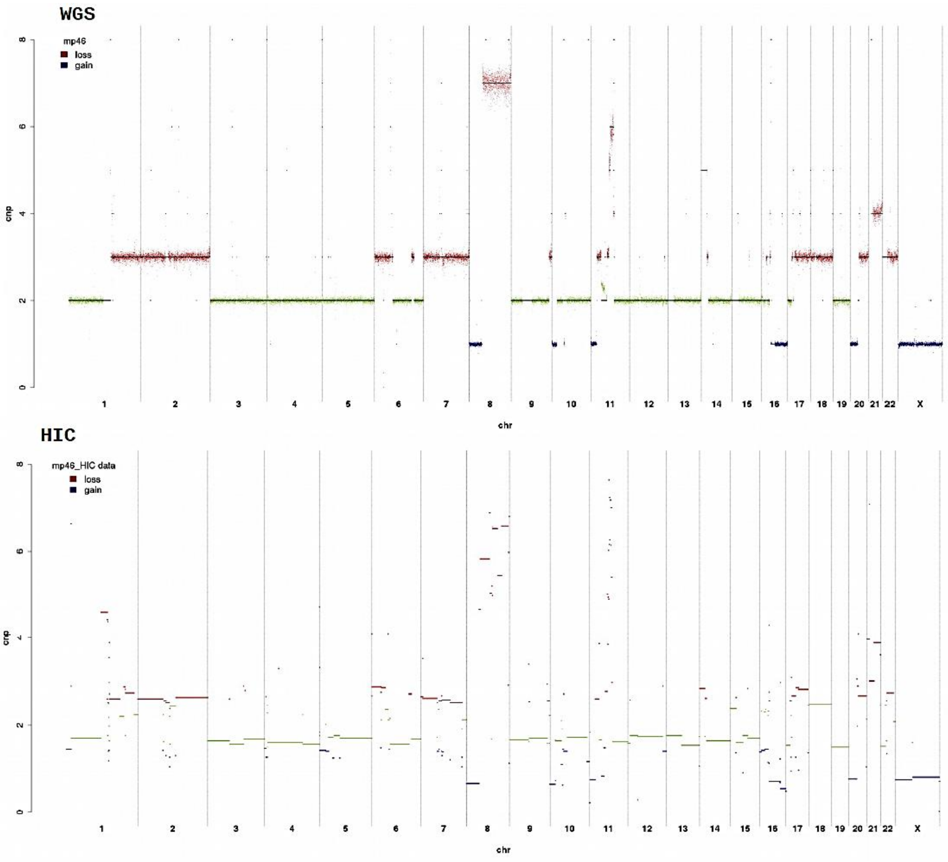

Supplementary Figure 9: Effect on compartment interaction scores after different normalization methods applied on Hi-C data generated for MP41, MP46 and NM.

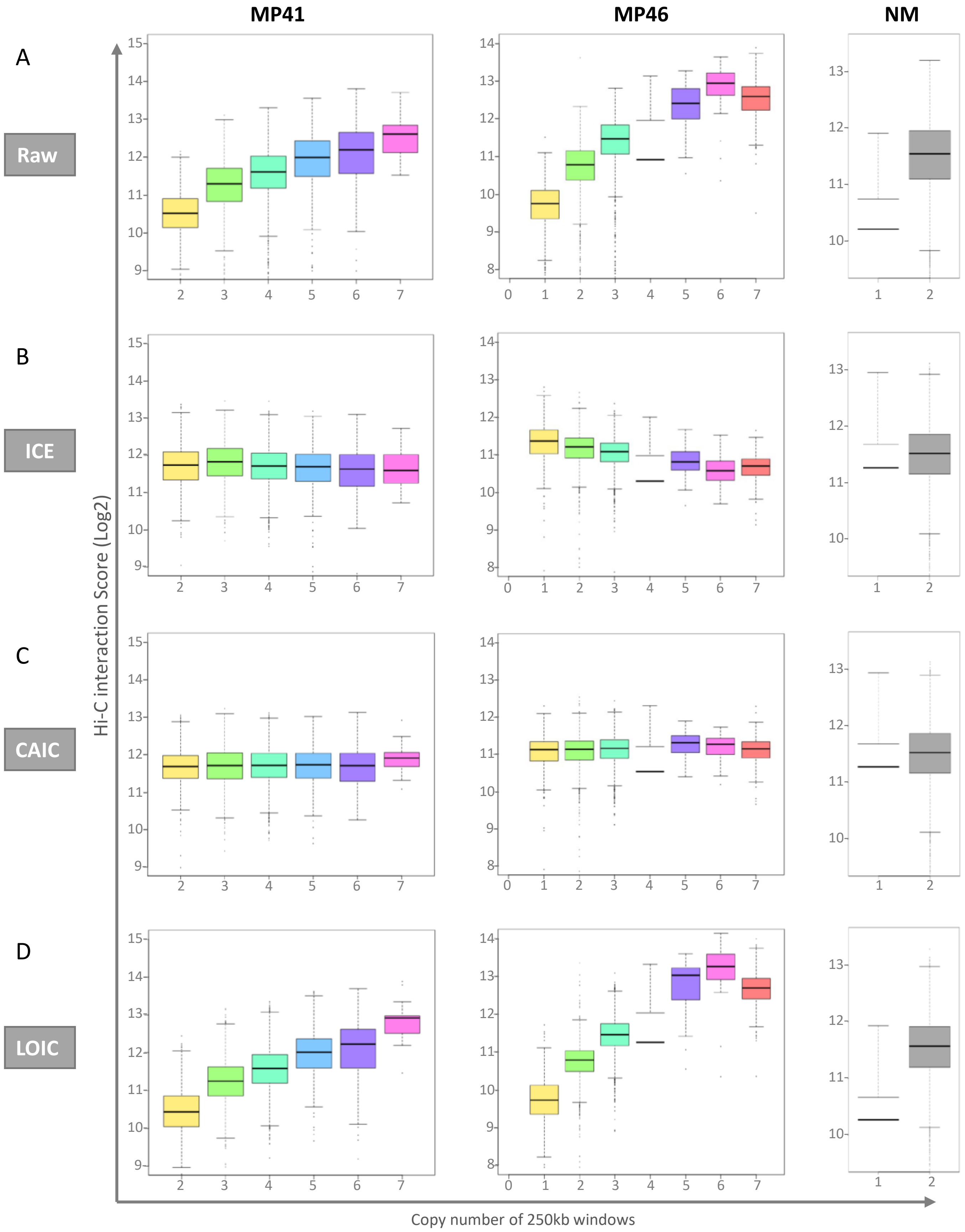

Supplementary Figure 10: Euchromatin and Heterochromatin compartments remain stable after CAIC normalization

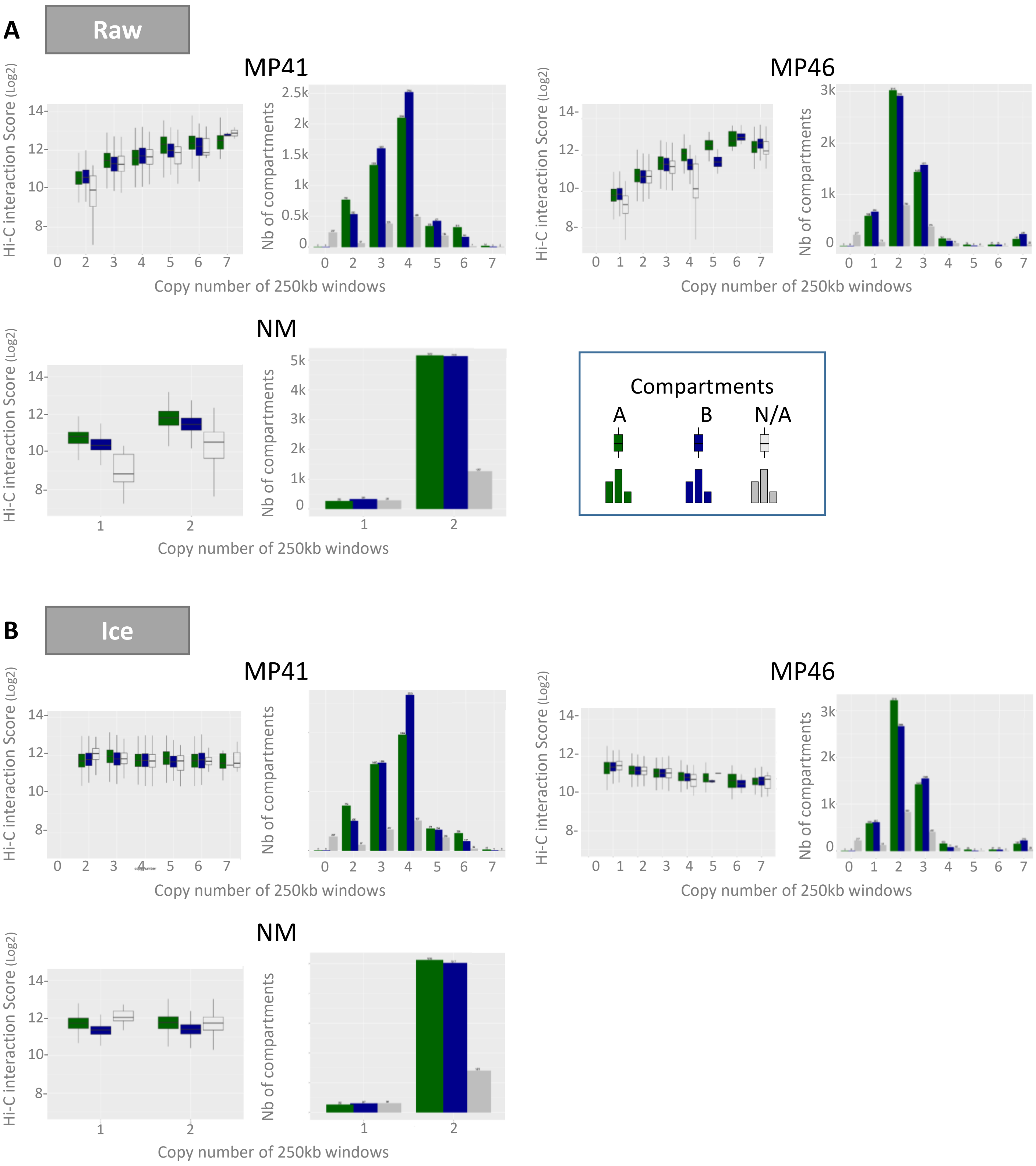

### Supplementary Figure 10: Euchromatin and Heterochromatin compartments remain stable after CAIC normalization

**C** CAIC

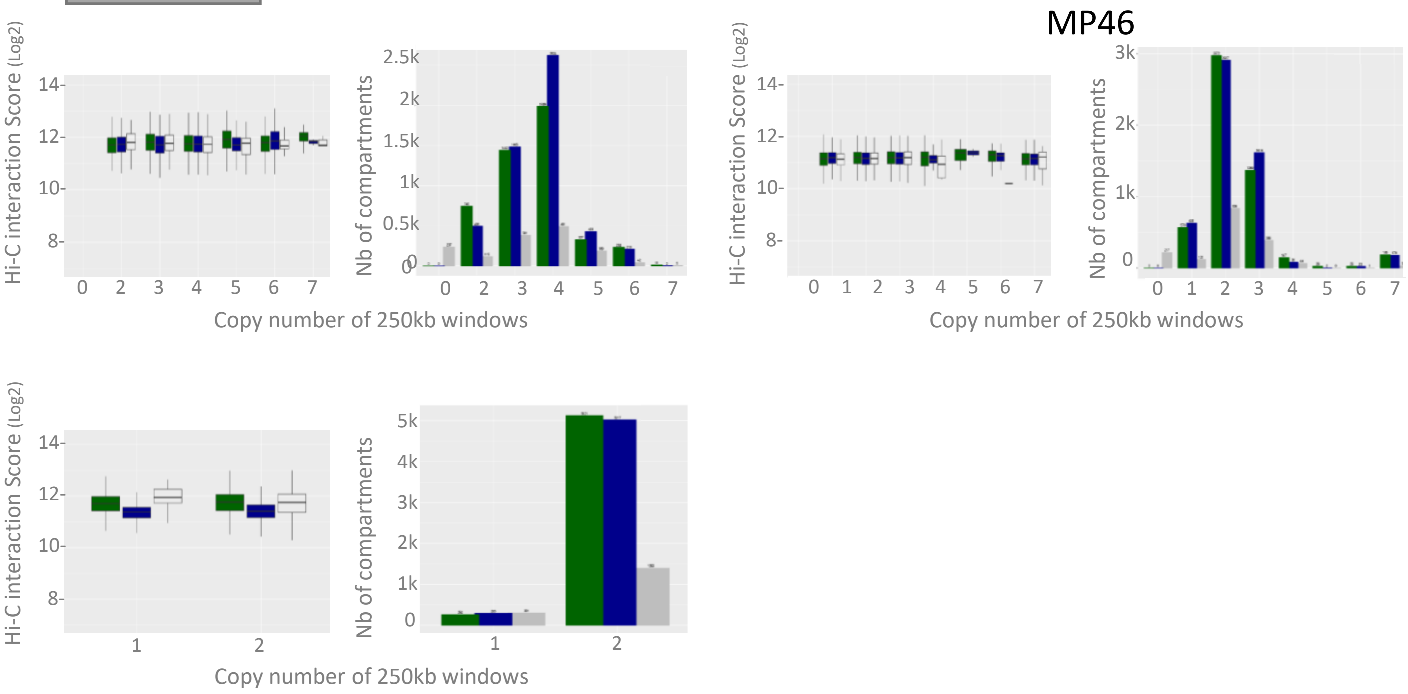


**D** LOIC

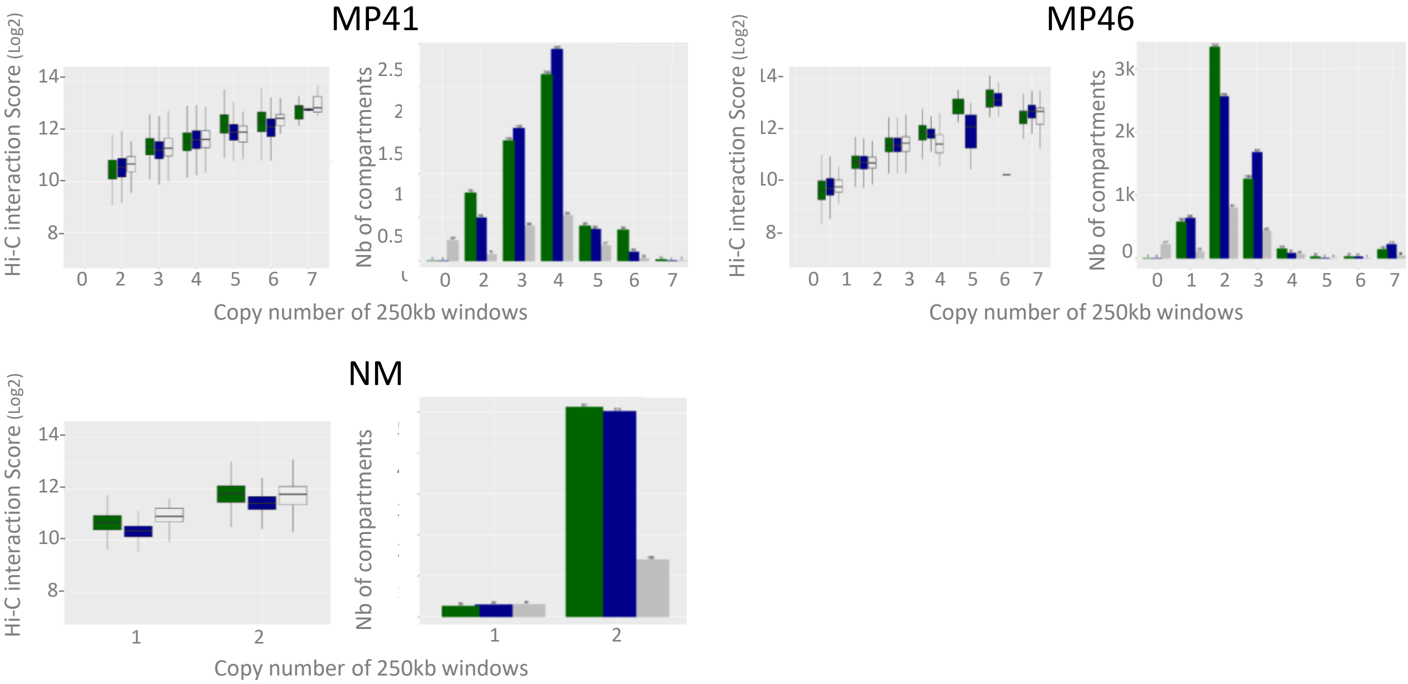

### Supplementary Figure 11: Prame transcription factors upregulated in UM models vs NM

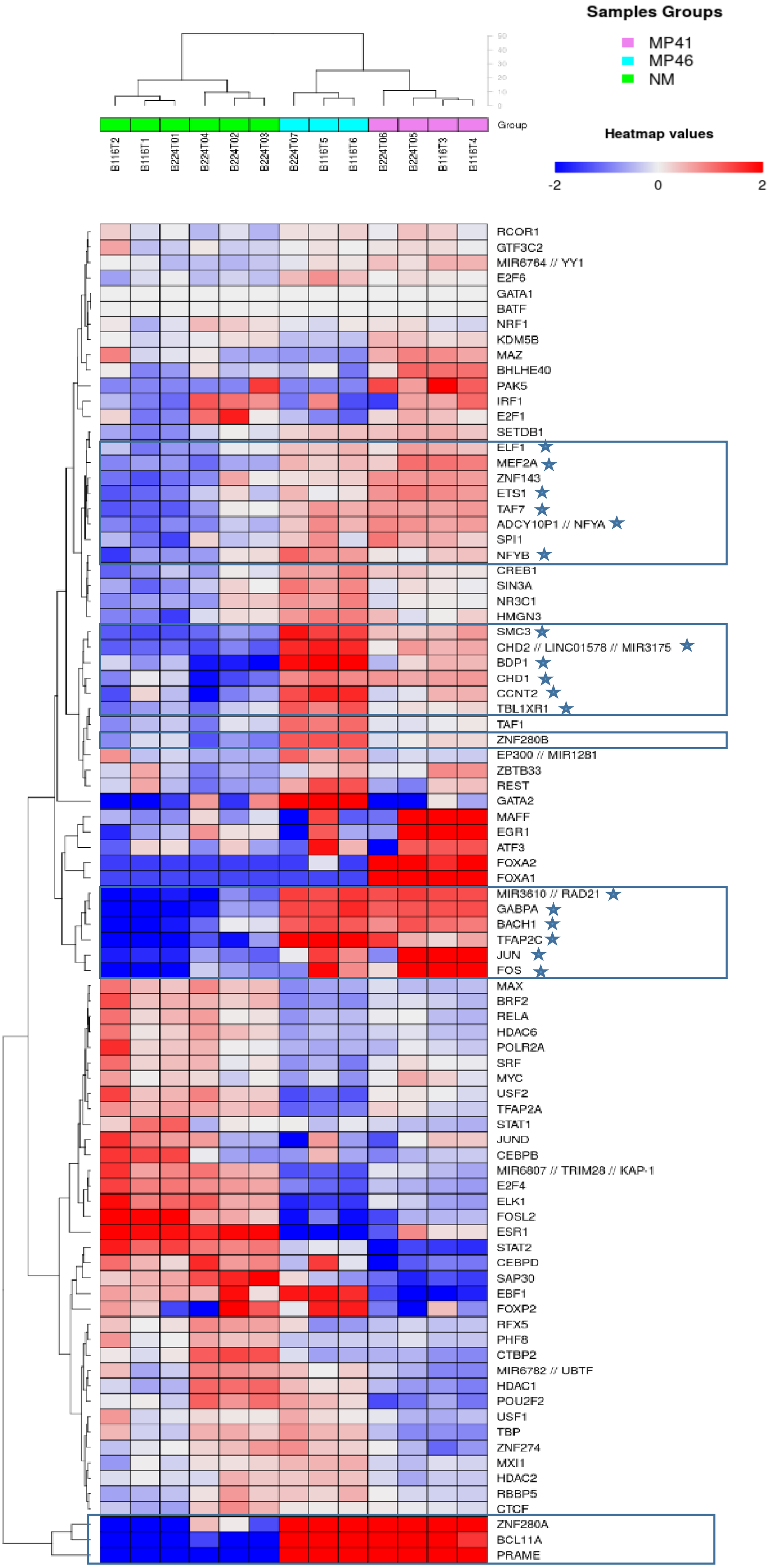
